# High throughput evaluation of genetic variants with prime editing sensor libraries

**DOI:** 10.1101/2022.10.26.513842

**Authors:** Samuel I. Gould, Alexandra N. Wuest, Kexin Dong, Grace A. Johnson, Alvin Hsu, Varun K. Narendra, Stuart S. Levine, David R. Liu, Francisco J. Sánchez Rivera

## Abstract

Many human diseases have a strong association with diverse types of genetic alterations. These diseases include cancer, in which tumor genomes often harbor a complex spectrum of single-nucleotide alterations and chromosomal rearrangements that can perturb gene function in ways that remain poorly understood. Some cancer-associated genes exhibit a tremendous degree of mutational heterogeneity, which may impact disease initiation, progression, and therapy responses. For example, *TP53*, the most frequently mutated gene in cancer, shows extensive allelic variation that leads to the generation of altered proteins that can produce functionally distinct phenotypes. Whether distinct variants of *TP53* and other genes encode proteins with loss-of-function, gain-of-function, or otherwise neomorphic phenotypes remains both controversial and technically challenging to assess, particularly at the endogenous level. Here, we present a high-throughput prime editing “sensor” strategy to quantitatively assess the functional impact of diverse types of endogenous genetic variants. We used this strategy to screen the largest collection of endogenous cancer-associated *TP53* variants assembled to date, identifying both known and novel alleles that impact p53 function in mechanistically diverse ways. Intriguingly, we find that certain types of endogenous *TP53* variants, particularly those in the p53 oligomerization domain, display opposite phenotypes in exogenous overexpression systems. These include disease-relevant variants found in humans with cancer predisposition syndromes that encode altered proteins with unique molecular properties. Our results emphasize the physiological importance of gene dosage in shaping native protein stoichiometry and protein-protein interactions, highlight the dangers of using exogenous overexpression systems to interpret pathogenic alleles, and establish a powerful computational and experimental framework for studying diverse types of genetic variants in their endogenous sequence context at scale.

## Introduction

A wide range of human diseases are associated with diverse genetic alterations that may be responsible for initiating, promoting, or otherwise modifying the course of a given disease. These alterations can be quite complex; for instance, cancer genomes typically contain a repertoire of single nucleotide variants (SNVs) and large-scale copy number alterations (CNAs) that can impact many genes in different ways depending on the type of alteration, gene function, and biological context. While tumor genotype is a well-established determinant of disease initiation, progression, and therapy responses, the functional impact conferred by the thousands of unique mutations observed in human tumors remains poorly understood. This presents a major challenge to precision medicine efforts that aim to tailor cancer therapies to patients suffering from cancers harboring specific genetic lesions. Beyond the clinic, understanding the impact that diverse types of mutations have on different residues and protein domains would improve our fundamental understanding of gene and protein function **(Fig. 1a)**.

**Fig. 1.**
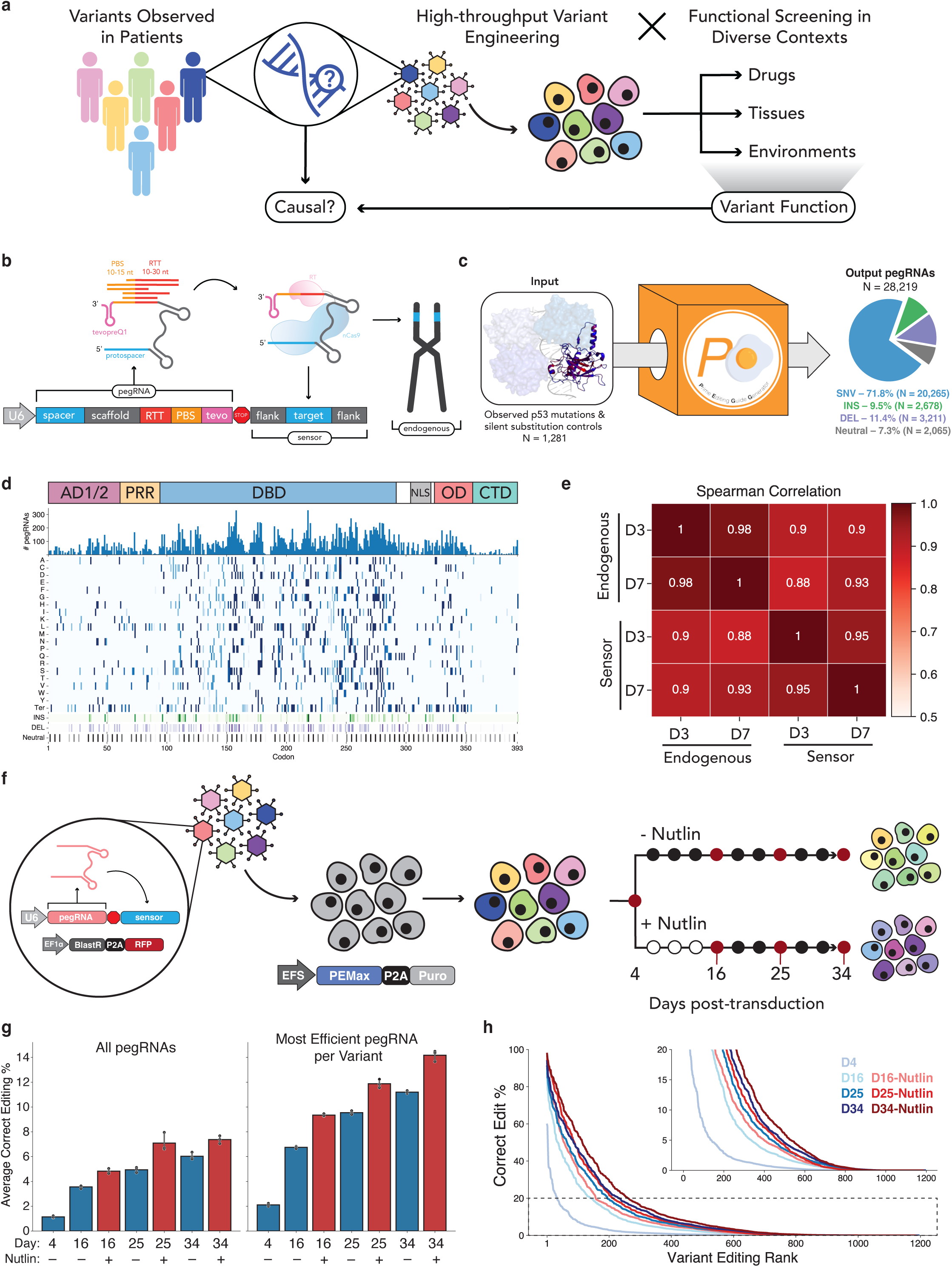
High Throughput design and evaluation of a *TP53* prime editing sensor library. **a,** Schematic of our overall approach. We aim to engineer variants observed in patients with high-throughput to perform functional screens in diverse contexts, elucidating variant functions to improve our ability to stratify and treat patients. **b,** Schematic of the sensor framework, which links each pegRNA to its editing outcome at the endogenous locus. RTT = reverse transcription template, PBS = primer binding site, tevo = tevopreQ1, RT = reverse transcriptase, nCas9 = nicking Cas9. **c,** We used PEGG to generate a *TP53* prime editing sensor library targeting over a thousand cancer-associated *TP53* variants with a median of 30 pegRNAs per variant. **d,** Heatmap visualization of the pegRNAs included in the *TP53* sensor library, which includes SNVs, indels, and silent substitutions. AD1/2 = activation domain 1/2, PRR = proline rich region, DBD = DNA binding domain, NLS = nuclear localization signal, OD = oligomerization domain, CTD = C-terminal domain. **e,** Correlation between editing at the sensor and endogenous locus in 8 *TP53*-targeting pegRNA-sensor pairs at Day 3 (D3) and Day 7 (D7) post-transduction. **f,** Schematic of the screening protocol. The prime editing sensor library is transduced into cells constitutively expressing PEmax, and screening is performed in the presence or absence of the small molecule Nutlin-3. **g,** The average correct editing percentage among all pegRNAs in the library (left) or when considering only the most efficient pegRNA for each variant (right) at various time-points in both conditions. **h,** Rank plot of the correct editing percentage of the most efficient pegRNA per variant, as assayed at the sensor locus, at each time-point.

Until recently, approaches for studying genetic variants have been limited to low-throughput, homology-directed repair (HDR) based methods or high-throughput, non-physiological gene overexpression systems^1–7^. While powerful, the former approach lacks scalability and generality due to the requirements of HDR and its limitation primarily to actively dividing cells. Gene overexpression systems have fewer requirements and are scalable but fail to physiologically recapitulate the biology driven by these variants due to the absence of endogenous gene regulation mechanisms, many of which are not known for genes of interest. The recent development of precision genome editing tools, including base editing and prime editing, allows variants to be modeled in their native endogenous genomic context with increased editing efficiency and theoretically higher throughput^8–10^.

Prime editing^10^ can be used to generate effectively any type of small mutation, including all SNVs and small insertions and deletions (indels). Prime editors are directed to engineer a mutation of interest by the instructions encoded in a prime editing guide RNA (pegRNA), which contains both a protospacer (the “search” sequence) and a 3’ extension sequence (the “replace” sequence that dictates the mutation installed at the site). The modular search-and-replace ability of prime editing has been leveraged to interrogate endogenous variants in high-throughput^11–13^. In these approaches, libraries of pegRNAs are transiently or stably delivered to cells expressing prime editors, and the fitness of variants is assessed by determining the relative distribution of endogenous alleles and/or pegRNAs. While powerful, these approaches have important limitations for screening applications, including 1) reliance on a small number of variant-specific pegRNAs with unknown editing performance, 2) inability to quantitatively assess endogenous genome editing at scale, and 3) potential overrepresentation of undesired indels due to using PE3.

With these challenges in mind, we sought to develop an integrative computational and experimental framework for high-throughput design, screening, and deconvolution of pegRNA libraries to interrogate a diverse spectrum of genetic variants. This includes pairing each pegRNA with a variant-specific synthetic “sensor” site^14^ that recapitulates the native architecture of the endogenous target locus. Doing so allows linkage of pegRNA identity to editing outcomes for simultaneous high-throughput quantification of pegRNA editing activity and empirical calibration of screening data.

We chose the p53 transcription factor as a prototype to test the utility of this approach for investigating the biological impact of specific genetic variants. Notably, *TP53* is the most frequently mutated gene in cancer and exhibits significant allelic variation, leading to the generation of altered proteins that can produce functionally distinct phenotypes. Whether distinct variants of *TP53* and other genes encode proteins with loss-of-function, gain-of-function, or otherwise neomorphic activities that influence cancer phenotypes is both controversial and technically challenging to investigate, particularly in high-throughput. Multiple studies have used cDNA-based exogenous overexpression systems to probe the fitness of p53 variants in human, mouse, and yeast systems^6,7,15,16^. However, given the artificial nature of these screens, which rely on expression of variants at supraphysiological levels, we hypothesized that these screens could have misrepresented one or more phenotypes associated with p53 variants. Artifacts that stem from exogenous overexpression systems could be particularly relevant when studying proteins like p53 because p53 functions as a tetramer whose expression and degradation is tightly controlled by the cell^17–19^. Thus, we reasoned that alterations to the stoichiometric balance of p53 via overexpression could lead to erroneous conclusions about the effects of particular p53 variants, including mis-classifying certain variants as non-causal or otherwise benign.

To tackle this question, we generated and screened a library of >28,000 pegRNAs targeting >1,000 *TP53* variants observed across >40,000 cancer patients^20^ — the largest set of endogenous *TP53* variants studied to date. We included SNVs, insertions, and deletions observed in patients, putative neutral silent substitutions as controls, and a panel of random indels to increase the functional search space. These experiments identified both known and novel alleles that impact p53 function in mechanistically diverse ways. Intriguingly, we discovered that certain types of endogenous variants, particularly those found in the p53 oligomerization domain, display opposite phenotypes when tested with exogenous overexpression systems. Collectively, these results highlight the physiological importance of gene dosage in shaping native protein stoichiometry and protein-protein interactions, and establish a powerful computational and experimental framework for studying diverse types of genetic variants at scale. To ensure widespread accessibility of this resource for the scientific community, we provide a publicly available Python package, Prime Editing Guide Generator (<u>pegg.readthedocs.io</u>), as a tool to generate prime editing sensor libraries.

## Results

### High-throughput design of prime editing sensor libraries with Prime Editing Guide Generator (PEGG)

A major limitation of using prime editing to systematically investigate genetic variants is the inherent variability in editing efficiency among different pegRNAs^10,21–23^. A number of computational tools for pegRNA design have been developed^24–33^, including machine learning algorithms that can nominate sets of pegRNAs predicted to produce high efficiency edits. However, even pegRNAs generated by these predictive algorithms require extensive experimental validation, and their editing activity is not guaranteed to correlate strongly across different cell types. We hypothesized that coupling pegRNAs with “sensors”—artificial copies of their endogenous target sites—would allow us to systematically identify high efficiency pegRNAs while controlling for the confounding effects of variable editing efficiency in a screening context **(Fig. 1b)**.

Synthetic sensor-like target sites have been previously used by our group and others to control for base editing gRNA editing efficiencies while defining the relative fitness of variants in genetic screens^14,34^. Multiple studies have applied a similar strategy to both base and prime editing technologies to identify features of efficient gRNAs or pegRNAs and train predictive algorithms^21,32,33,35–37^. However, this approach has yet to be applied for high-throughput phenotypic screening of endogenous genetic variants with prime editing, likely due in part to the lower editing efficiency of prime editing relative to base editing. We reasoned that a sensor-based prime editing screening approach could be powerful to discriminate *bona fide* endogenous variants from undesired editing outcomes that enrich or deplete in a screen. Moreover, the sensor approach would theoretically overcome the limitations of assessing variants at different genetic sites in parallel by eliminating the need to sequence multiple endogenous loci.

To test this approach, we first needed to build a computational tool capable of designing and ranking pegRNAs for thousands of genetic variants, while automatically generating a paired sensor site. To address this unmet need, we built and publicly released the Prime Editing Guide Generator (PEGG) **(Extended Data Fig. 1a)**, a Python package that enables high throughput design of prime editing sensor libraries^38^ (available at <u>pegg.readthedocs.io</u>). PEGG is compatible with a range of mutation input formats, including all of the datasets on the cBioPortal, ClinVar identifiers, and custom mutation inputs^39,40^.

We chose the *TP53* tumor suppressor gene as a prototype to establish and credential a scalable prime editing sensor-based screening approach for a number of reasons. First, *TP53* is the most frequently mutated gene in human cancer, with ∼50% of patients suffering from tumors harboring a mutation within the *TP53* gene while the rest often inactivate the p53 pathway through other mechanisms. Secondly, thousands of unique *TP53* mutations have been identified in cancer patients, including eight or so “hotspot” alleles in specific residues that exhibit the highest mutational frequencies^19^. Although p53 has been studied for decades, there have been few systematic studies, and those have been hampered by reliance on artificial overexpression of mutant p53 proteins, unrepresentative cell lines, and/or a limited spectrum of mutations evaluated^6,7,15,16^. These and other studies have sparked controversy in the field over whether any mutant p53 proteins are endowed with activities that go beyond loss-of-function or dominant negative activity to achieve gain-of-function or neomorphic status. These are important questions that extend beyond *TP53* because mutant gain-of-function proteins generated by cancer-associated variants, and the phenotypes they produce, could represent attractive therapeutic targets. Lastly, prime editing sensor-based screening could be scaled up and broadly deployed to identify causal genetic variants implicated in cancer and other diseases with a strong genetic association.

With the above goals in mind, we first sought to generate a library of pegRNAs targeting *TP53* variants. To generate this library, we selected variants from the MSK-IMPACT database, which uses deep exon sequencing of patient tumor samples to identify cancer-associated variants^20^. From this database of over 40,000 patients, we chose all observed SNVs in p53, as well as frequently observed insertions and deletions, along with a collection of random indels **(Extended Data Fig. 1b)**. We reasoned that including multiple pegRNAs with different protospacers and combinations of pegRNA properties for each variant would allow us to scan the pegRNA design space more thoroughly to identify highly efficient guides for robust statistical analysis of variant phenotypes. To accomplish this, we used PEGG to produce 30 pegRNA designs per variant (for pegRNAs with a sufficient number of accessible PAM sequences) with varying RTT (10–30 nt) and PBS lengths (10–15 nt) coupled to canonical “NGG” protospacers. The generated pegRNA designs were ranked based on a composite “PEGG score” that integrates literature best practices for pegRNA design **(Extended Data Fig. 1a, Supplemental Table 1)**.

PEGG also generated silent substitution variants as neutral internal controls for the screen. We also filtered pegRNAs to exclude protospacers with an MIT specificity score below 50 to reduce the probability of off-target editing^41^ **(Extended Data Fig. 1e)**. In addition, these pegRNA designs included an epegRNA motif, tevopreQ_1_, an RNA pseudoknot located at the 3′ end of the pegRNA that improves editing by preventing degradation of the guide^22^. Even after these relatively stringent filtration steps, we were able to generate pegRNA designs for more than 95% of the input variants, resulting in a library of >28,000 pegRNAs **(Fig. 1c, 1d; Extended Data Fig. 1c, 1d)**. Each pegRNA in the library is also paired with a 60-nucleotide long variant-specific synthetic “sensor” that is generated by PEGG and included in the final oligonucleotide design. Every sensor is designed to recapitulate the native endogenous target locus, thereby linking pegRNA identity to editing outcomes **(Fig. 1b)**.

To test the efficacy of using the sensor as a readout of editing at the endogenous locus, we randomly selected eight *TP53* variant-specific pegRNA sensors generated during the process of library preparation. We generated lentivirus for each of these PE sensor constructs and performed separate transductions into cells expressing PEmax. At 3- and 7-days post-transduction, we harvested genomic DNA and amplified both the pegRNA–sensor cassette and the endogenous locus targeted by each pegRNA. Analysis of editing at the sensor and endogenous locus revealed a very high correlation between the sensors and endogenous sites (Spearman correlation ≥ 0.9; **Fig. 1e**). In general, the prime editing sensor seems to slightly overestimate the editing activity at the endogenous locus, likely in part due to differences in locus chromatin accessibility^42^, but the ranking of pegRNA editing efficiencies is largely preserved, validating our sensor-based approach.

### High-throughput interrogation of *TP53* variants using prime editing sensor libraries

Next, we screened these variants in *TP53* WT A549 lung adenocarcinoma cells stably expressing PEmax^21^. To measure prime editing activity, we generated and transduced these cells with a modified all-in-one lentiviral version of the fluorescence-based PEAR reporter^43^, which displayed strong editing activity **(Extended Data Fig. 2a)**. We then introduced the lentiviral *TP53* sensor library into these cells at a low multiplicity of infection and in triplicate while ensuring a library representation of >1000X at every step of the screen. Four days post-transduction, we split the populations into untreated or Nutlin-3-treatment arms **(Fig. 1f)**. Nutlin-3 is a small molecule that inhibits MDM2 to activate the p53 pathway, and some *TP53* mutations have been shown to promote bypass of p53-dependent cell cycle arrest and apoptosis^44^. We hypothesized that this treatment group may increase the signal-to-noise ratio between *TP53* variants with loss-of-function (LOF) and gain-of-function (GOF) activities. We allowed the screen to progress for 34 days, harvesting cell pellets from each replicate and treatment arm at multiple time-points **(Extended Data Fig. 2b)**. Genomic DNA extracted from each sample was used to amplify the pegRNA–sensor cassettes, which were subjected to next-generation sequencing to simultaneously assess enrichment/depletion of pegRNAs and their editing activity and outcomes at the sensor target site **(Extended Data Fig. 2c, 2d)**.

The average editing efficiency among all pegRNAs in the library increased in a time-dependent manner, peaking at ∼8% in the final time-point. In general, we observed low indel rates and strong correlation in sensor editing among replicates **(Fig. 1g, Extended Data Fig. 3a-d)**. Strikingly, selecting only the most efficient pegRNA design for each variant led to a two-fold increase in the average editing efficiency, highlighting the utility of the sensor for systematic empirical identification of high efficiency pegRNAs **(Fig. 1g, Extended Data Fig. 3e-g).** Cells with higher editing efficiency also exhibited stronger Nutlin-3 bypass in the Nutlin-3-treatment arm **(Fig. 1g).** Based on the assessment of editing at the sensor locus, we were able to identify active pegRNAs (≥2% editing efficiency) for more than half of the *TP53* variants included in the library. This includes highly efficient pegRNAs that install the desired edit with over 20% efficiency for more than 20% of the variants **(Fig. 1h)**. These pre- validated pegRNAs could be further engineered with silent mutations that evade mismatch repair to boost overall editing efficiency^21^.

The size and diversity of this library also allowed us to examine features of highly efficient pegRNAs, which broadly recapitulated previous observations^32,33,37,45^. Correlation analysis between various pegRNA features and editing efficiency across all time-points identified the estimated on-target activity of the protospacer (as predicted by Rule Set 2)^46^ as the single largest determinant of prime editing efficiency **(Fig. 2a).** In addition, the distance between the edit and the nick introduced by nCas9 was negatively correlated with editing efficiency, while the length of the post-edit homology was positively correlated with editing efficiency **(Fig. 2a).** Thus, edits closer to the nick and with larger post-edit homology were more efficient, consistent with previous findings^32,33,37,45^.

**Fig. 2.**
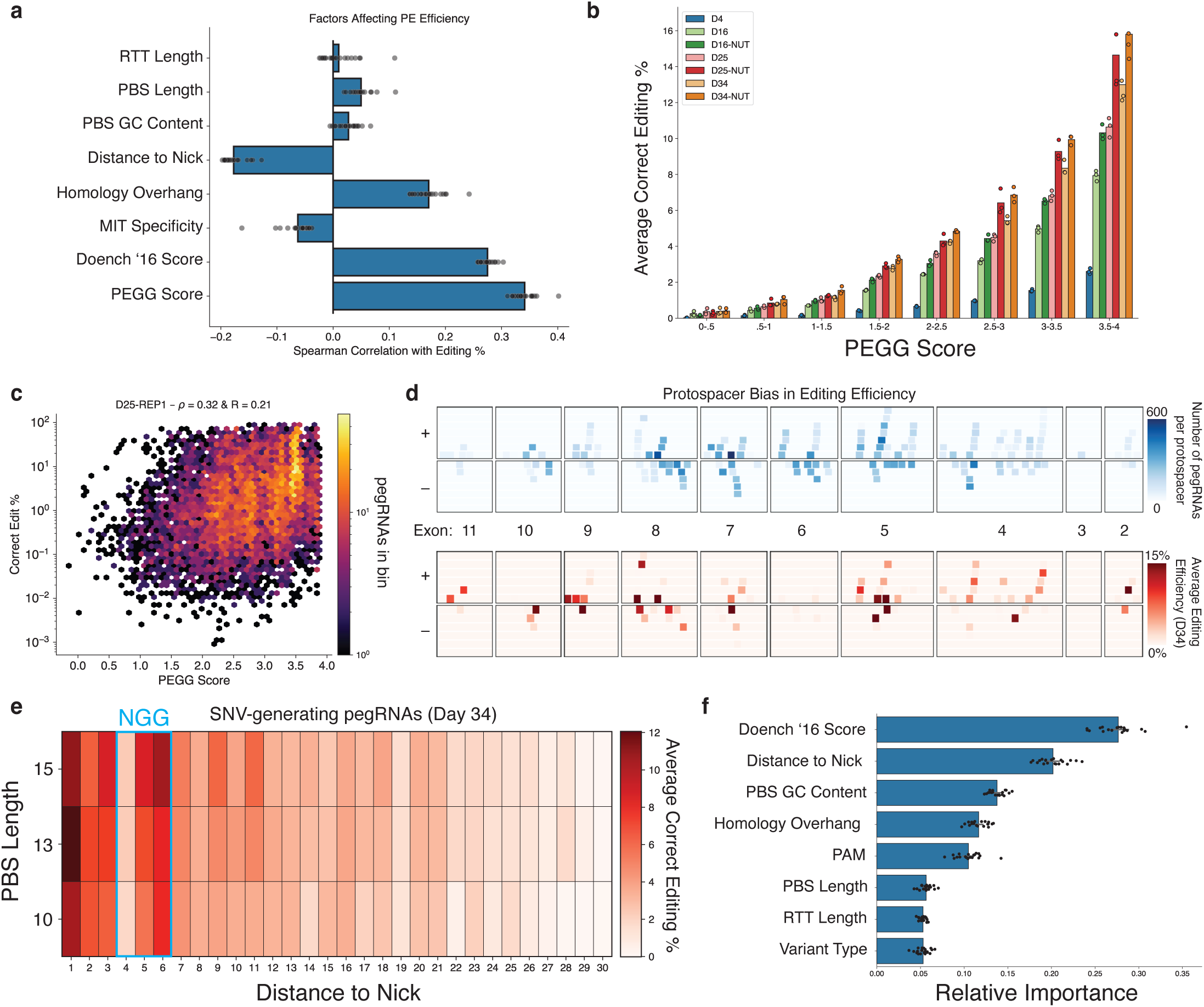
Identification of features of highly efficient pegRNAs. **a,** Spearman correlations between various features of pegRNA design and correct editing percentage, assessed for all pegRNA-sensor pairs with sufficient reads. Each dot represents a separate replicate/time-point. **b,** The relationship between PEGG score and average correct editing percentage at each time-point & condition is monotonically increasing. **c,** A representative example of the correlation between PEGG score and editing efficiency for Day-25 Replicate 1 (Untreated). Spearman correlation = 0.32; Pearson correlation = 0.21. **d,** Visualization of the protospacer bias in editing efficiency. The number of pegRNAs generated per protospacer at each TP53 exon on the plus (+) or minus (-) strand (top) and the average editing efficiency at each of these protospacers at Day 34 of the untreated condition (bottom). **e,** Average editing efficiency for SNV-generating pegRNAs in the library as a function of distance to the nick generated by PEmax and PBS length. The location of the "NGG" PAM sequence is highlighted in blue. Protospacer disrupting (locations +1 to +3) and PAM-disrupting variants (locations +5 & +6) tend to be more efficient. **f,** Feature importances of 20 Random Forest models separately trained to predict pegRNA efficiency. Each dot represents a different model.

Notably, the PEGG Score, which is a weighted linear combination of pegRNA features based on literature best practices, correlated more strongly with prime editing efficiency than any other single feature, achieving a Spearman correlation of up to 0.4 **(Fig. 2a-c).** Though this correlation is modest relative to published predictive models^32,33,37^, the PEGG score is a simple, unbiased, and cell type/organism-agnostic prediction of pegRNA activity that could complement machine learning-based predictions of PE activity, which may vary due to training on particular cell types.

To further analyze the differences in prime editing activity among the 173 protospacers spanning the *TP53* locus, we visualized the number of pegRNAs that utilized each protospacer and the average editing efficiency at each protospacer **(Fig. 2d)**. This analysis suggests that only a subset of protospacers can be used to generate high efficiency pegRNAs, while other protospacers retain little to no editing activity. We also found that pegRNAs that introduce edits that disrupt the protospacer or PAM sequence tend to be more efficient **(Fig. 2e)**. Relative to the nick created by nCas9, SNVs introduced at the +1–3 position, which mutate the protospacer, and at the +5–6 position, which mutate the guanine bases in the NGG PAM, display increased editing activity. In contrast, edits introduced at the +4 position, corresponding to the “N” in the “NGG” PAM sequence, display reduced editing likely due to their failure to disrupt the PAM sequence **(Fig. 2e).**

Finally, we trained a random forest regressor to predict pegRNA efficiency **(Extended Data Fig. 4a).** Even with a restricted set of features, this algorithm was able to predict pegRNA activity with a Spearman correlation of ∼0.6, comparable to other, more complex algorithms that are used to predict PE activity^32,33,37^ **(Extended Data Fig. 4b)**. Analysis of the relative feature importances of this random forest model was again consistent with previous findings, and highlighted the GC content of the PBS as another important determinant of editing not identified with simple correlation analysis **(Fig. 2f).** These results demonstrate that large-scale, gene-specific prime editing sensor screening datasets can also provide insight into the determinants of high efficiency prime editing, even though these libraries were not designed with that objective in mind.

### Sensor-based calibration of prime editing screening data identifies pathogenic *TP53* variants

To assess the relative fitness conferred by engineered *TP53* variants, we used the MAGeCK pipeline to normalize read counts among replicates and quantify the log^2^ fold-change (LFC) and false discovery rate (FDR) of pegRNAs in the library^47^. While the LFC in pegRNA counts was highly correlated in replicates from the untreated and Nutlin-3-treated arms of the screen, respectively, the correlation among replicates between the two conditions was modest, suggesting that treatment-dependent biological effects were occurring **(Extended Data Fig. 3a, 5).** To investigate this further, we used the sensor target site as a quantitative proxy for editing efficiency at the endogenous locus to systematically filter pegRNAs based on their empirical editing efficiency **(Fig. 3a-d, Extended Data Fig. 6a-d)**. As expected, the number of significantly enriched or depleted pegRNAs in “sensor-calibrated” datasets was generally inversely proportional to their editing efficiency, prompting us to focus our statistical analyses on a dataset composed of pegRNAs with ≥10% editing efficiency **(Fig. 3a-e)**. These results demonstrate that our sensor-based approach allows empirical removal of pegRNAs that exhibit potentially spurious enrichment or depletion, and low and/or undesired editing activity, retaining pegRNAs that are significantly more likely to introduce the variants of interest.

**Fig. 3.**
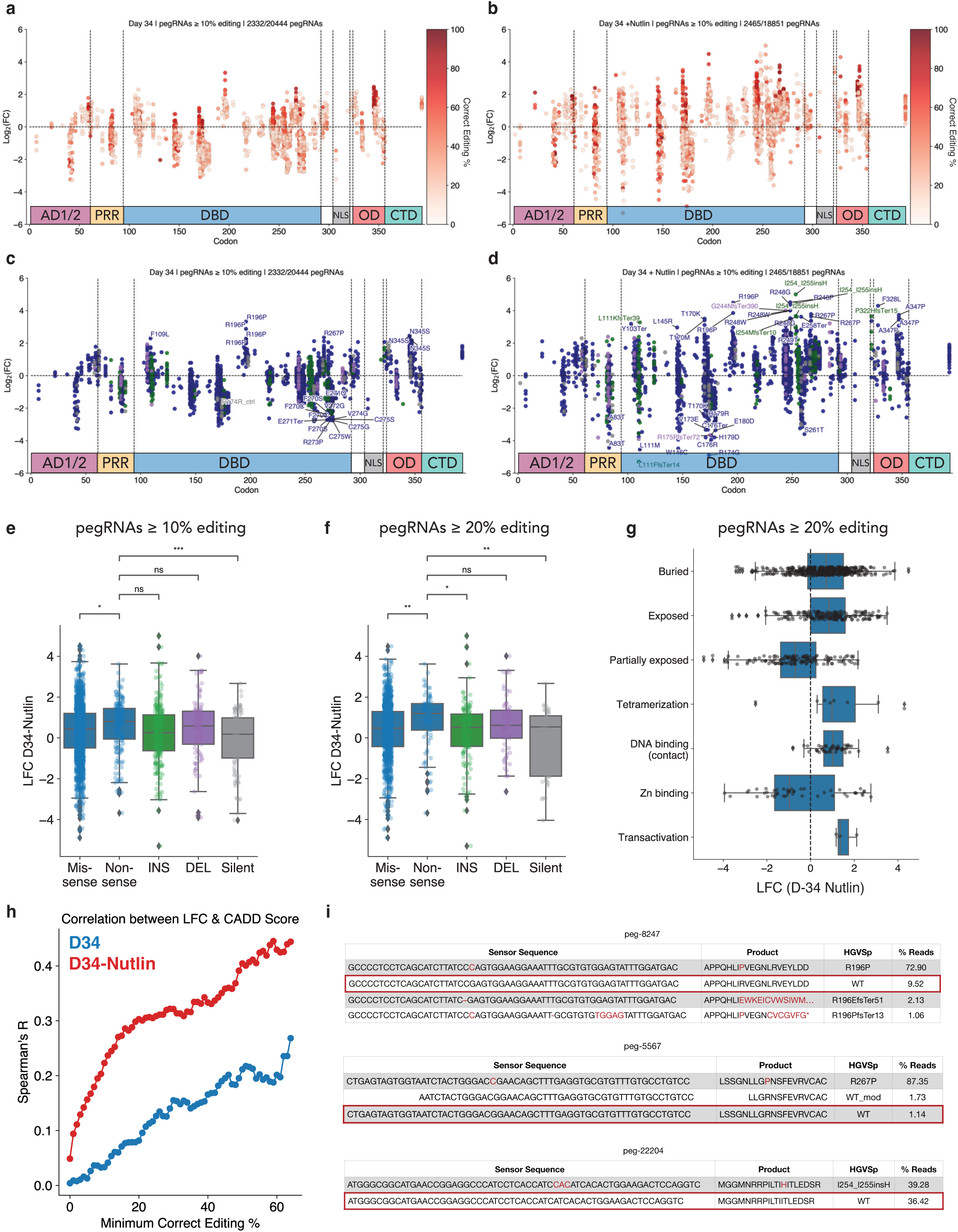
High-throughput prime editing sensor screens identify pathogenic *TP53* variants. **a,** The LFC of each pegRNA 2: 10% editing with at least 10 sensor reads at Day 34 relative to Day 4 in the untreated condition, with pegRNAs colored by editing efficiency. AD1/2 = activation domain 1/2, PRR = proline rich region, DBD = DNA binding domain, NLS = nuclear localization signal, OD = oligomerization domain, CTD = C-terminal domain. **b,** The LFC of each pegRNA 2: 10% editing with at least 10 sensor reads at Day 34 relative to Day 4 in the Nutlin-treated condition, with pegRNAs colored by editing efficiency. **c,** Plot (a), but with pegRNAs colored by variant type. Enriching pegRNAs with LFC 2: 2 and FDR < .05 labeled. Depleting pegRNAs with FDR < .05 labeled. Blue = SNV, Green = INS, Purple = DEL, Gray = Silent. **d,** Plot (b), but with pegRNAs colored by variant type. Selected enriching pegRNAs with FDR < .05 labeled. Depleting pegRNAs with FDR < .05 labeled. Blue = SNV, Green = INS, Purple = DEL, Gray = Silent. **e, f,** Box-plot of LFC in pegRNAs at Day 34 in Nutlin-treated condition separated by variant class for pegRNAs 2: 10% editing and 2: 20% editing. Statistics shown for t-test of independent samples with Bonferroni correction. * = p-value ::5 .05, ** = p-value ::5 .01, *** = p-value ::5 .001, **** = p-value ::5 .0001, ns = not significant (p-value > .05). **g,** Box-plot of the LFC at Day 34 in the Nutlin-treated condition for pegRNAs 2: 20% editing with annotated residue functions. **h,** Spearman correlation between LFC of SNV-generating variants and CADD score at Day 34 in untreated (blue) and Nutlin-treated (red) conditions at minimum correct editing thresholds spanning 0% to 65%. **i,** Sensor editing for selected pegRNAs at Day 34 in the Nutlin-treated condition.

As hypothesized, the dynamic range in the Nutlin-3-treated arm of the screen was significantly higher than the untreated arm, with pegRNAs more strongly enriching and depleting in the presence of Nutlin-3 **(Fig. 3a, 3b).** Treatment with Nutlin-3 also selected for cells with higher efficiency editing, improving the resolution of the screen **(Fig. 3a, 3b)**. Several putative pathogenic *TP53* variants, including R196P and R267P, were strongly enriched in both treatment arms, with multiple pegRNAs per variant appearing as top hits **(Fig. 3c, 3d).** Given the higher dynamic range of the Nutlin-3 treatment arm, as well as the possibility that this treatment biases towards the discovery of dominant negative *TP53* mutations^2,6,16^, we focused our analyses on this treatment group. Multiple *TP53* variants showed significant enrichment in the Nutlin-3-treatment group, including SNVs and indels in the C-terminal half of the DNA-binding domain (DBD) and the oligomerization domain (OD) **(Fig. 3d)**. This includes multiple variants at residues 248 and 249, which are known mutational hotspots in p53 and commonly observed in cancer patients and individuals with the Li-Fraumeni cancer predisposition syndrome^17^.

Provocatively, the most commonly observed *TP53* mutation, R175H, did not show strong enrichment despite the existence of multiple R175H pegRNAs exhibiting ≥10% editing efficiency. In fact, the majority of the top enriching variants we identified were not in known mutational hotspots^19^, suggesting that other types of variants can produce stronger phenotypes. These include the top hit, an insertion of a histidine between residues 254 and 255, as well as R196P and multiple variants in the oligomerization domain (F328L, N345S, A347P) **(Fig. 3d).** These observations are consistent with the possibility that some *TP53* hotspot mutations are observed in part due to disproportionately high mutagenesis rates at the genomic level due to extrinsic and intrinsic factors, such as tobacco smoke and APOBEC activity, rather than only due to the fitness advantage conferred by these variants relative to other *TP53* mutations^6,19^. We also identified strongly depleting variants in the DBD that may retain wild type p53 transcriptional activity or fail to exert a dominant negative effect on the p53 tetramer **(Fig. 3d).** Collectively, these results validate the utility of our approach and dataset to accurately identify pathogenic *TP53* variants.

Bulk quantification of pegRNAs grouped by variant class also revealed that nonsense variants were significantly more enriched compared to missense and silent variants. This is evident at multiple thresholds of pegRNA activity **(Fig. 3e, 3f).** As expected, silent variants tend to deplete, particularly when considering pegRNAs at higher threshold for editing efficiency (≥ 20%), bolstering our confidence in the fidelity of the screen **(Fig. 3f).** Using available annotations of p53 residue function^48^, we also found that, as expected, variants in DNA binding/contacting residues displayed strong enrichment (including residues 248 and 273) **(Fig. 3g)**. Intriguingly, certain variants involved in tetramerization and transactivation (e.g., L22V) were also strongly enriched, despite the low frequency of mutation in these residues **(Fig. 3g)**. Other observations are more difficult to interpret, such as the large variance in the enrichment of mutations that affect residues involved in zinc binding, or the fact that variants in partially exposed residues tended to deplete while those in buried and exposed residues tended to enrich **(Fig. 3g).** Altogether, these observations suggest that there is a large, underappreciated phenotypic variance in the relative fitness conferred by distinct *TP53* variants — not all p53 variants are one and the same, a concept that is likely relevant across many other genes and more broadly in biology.

We also sought to quantify the degree of concordance between our screening results and widely used metrics of variant deleteriousness. To do so, we used the Combined Annotation-Dependent Depletion (CADD) score, which integrates evolutionary conservation of residues with other metrics of pathogenicity to generate a CADD score, with higher scoring variants predicted to be more deleterious^49^. We observed a low correlation between CADD score and enrichment of all SNV-specific pegRNAs. However, the correlation between CADD score and variant-specific fitness increased dramatically when we used sensor target sites to restrict our analysis to variants generated by high efficiency pegRNAs **(Fig. 3h)**. We achieved a Spearman correlation of ∼0.3 when considering Nutlin-3-treated pegRNAs with ≥ 15% editing activity, and >0.4 when considering pegRNAs ≥ 50% editing **(Fig. 3h)**. Across all minimum pegRNA editing activity thresholds, the CADD score correlated more strongly with fitness of Nutlin-3-treated pegRNAs to the untreated condition. These results emphasize the significant advantage of including sensor target sites in prime editing screening libraries to assess pegRNA efficiency and reinforce the ability of Nutlin-3 to effectively pull out genuine LOF, GOF, and separation-of-function *TP53* variants **(Fig. 3h, 3i).**

### Sequencing of endogenous *TP53* validates the prime editing sensor approach

The above results demonstrate that sensor-calibrated quantification of pegRNA enrichment and depletion can be used to identify *bona fide* pathogenic *TP53* variants. However, these analyses do not rule out the possibility that these changes are independent of true editing at the endogenous *TP53* locus. To formally test if prime editing sensor screens can faithfully quantify the effects of variants engineered at endogenous loci, we performed targeted next-generation sequencing of specific regions in exons 6, 7, and 10 of *TP53* using genomic DNA extracted from untreated and Nutlin-treated cells from Day 4 and Day 34 timepoints **(Fig. 4a)**. We reasoned that sequencing the native *TP53* locus would allow us to directly compare the fold-change in pegRNA counts with the fold-change of variants installed at defined targeted sites within endogenous *TP53*.

**Fig. 4.**
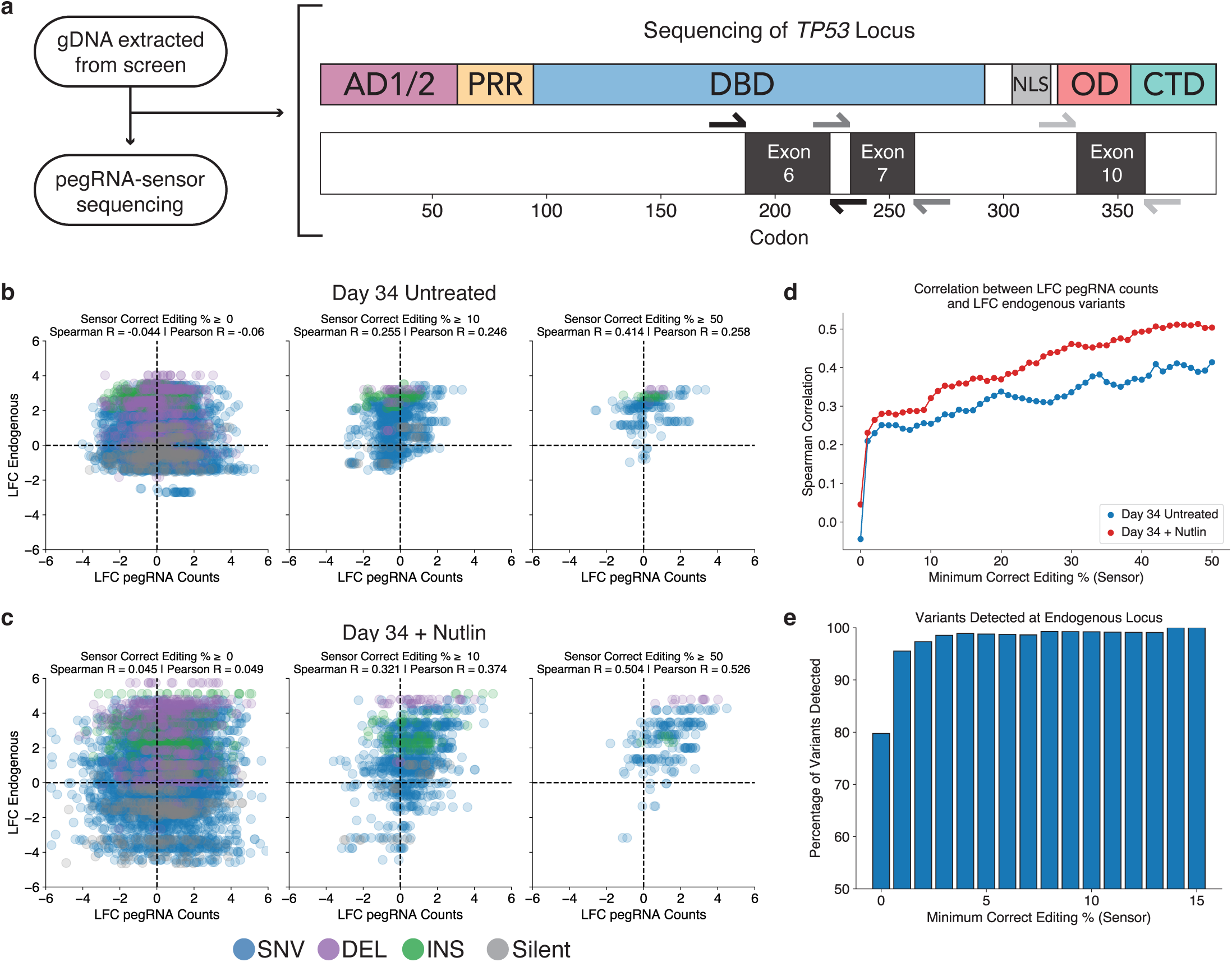
Sequencing of TP53 validates the prime editing sensor approach. **a,** Schematic of the sequencing of the TP53 locus. Regions within exons 6, 7, and 10 were sequenced from genomic DNA extracted at Day 4 and Day 34 in the untreated and Nutlin-treated arm of the screen. **b,** Correlation between the LFC in pegRNA counts and the LFC of endogenous variants at the TP53 locus for Day 34 vs. Day 4 of the untreated arm of the screen at different thresholds of pegRNA editing activity. **c,** Correlation between the LFC in pegRNA counts and the LFC of endogenous variants at the TP53 locus for Day 34 vs. Day 4 of the Nutlin-treated arm of the screen at different thresholds of pegRNA editing activity. **d,** Spearman correlation between the LFC in pegRNA counts and the LFC of endogenous variants at the TP53 locus as a function of the minimum sensor correct editing threshold for Day 34 untreated (blue) and D34 Nutlin-treated (red) samples. **e,** The percentage of variants detected (2 500 counts) at the TP53 locus at Day 34 (untreated) for variants with pegRNAs at different minimum sensor correct editing thresholds.

Unlike non-sensor-based prime editing screens, which are likely to suffer from significant noise during enrichment due to the difficulty of designing high efficiency pegRNAs, our sensor-based prime editing screen could be empirically denoised by filtering for pegRNAs that edited their cognate sensor above a given editing threshold. Indeed, we observed no correlation between the LFC in pegRNA counts with the corresponding LFC of endogenous variants engineered in the native *TP53* locus when considering all pegRNAs **(Fig. 4b-d)**. However, taking advantage of the sensor site to filter out pegRNAs below a given correct editing threshold dramatically reduced the noise in this data and revealed a strong correlation between the fold-change in pegRNA counts and endogenous variant counts **(Fig. 4b, 4c)**. This correlation increased monotonically as the minimum correct editing threshold increased, reaching a Spearman correlation > 0.4 in the untreated arm and > 0.5 in the Nutlin-treated arm of the screen when using a minimum correct editing threshold of 50% **(Fig. 4d)**. Moreover, edited exonic sites targeted by top enriching pegRNAs (e.g. R196P, A347P, I254_I255insH) were significantly enriched relative to their wild type counterparts and strongly correlated with their respective pegRNA counts. In addition, we were able to detect nearly all of the variants engineered by active pegRNAs (≥ 500 counts per variant and ≥ 1% correct sensor editing), and the detectable fraction reached saturation when considering pegRNAs producing ≥ 14% editing at the sensor site **(Fig. 4e)**. Altogether, these results emphasize the essentiality of integrating quantitative, sensor-like approaches to accurately extract true signal from the high levels of noise that are inherent in large-scale prime editing screens. Indeed, our analysis demonstrates that screening pegRNAs without any empirical quantification of their editing activity invariably leads to spurious conclusions concerning the fitness of the variants that those pegRNAs are intended to engineer.

### Functional validation of pathogenic *TP53* variants identified with prime editing screening

The above data demonstrate that pegRNA-specific sensor modules can be used to rigorously calibrate screening results to limit the analysis of variant fitness effects only to highly efficient pegRNAs. Though suggestive, these results do not formally prove that top scoring pegRNAs are enriched due to the introduction of defined genetic variants at the endogenous target locus and that these drive the observed biological differences. To test this, we selected a cohort of 29 pegRNAs that significantly enriched or depleted in the screen, or that generated commonly observed “hotspot” mutations **(Fig. 5a, 5b)**. This set of pegRNAs targeted residues that spanned the *TP53* locus and also included two control pegRNAs that install silent edits **(Fig. 5a, 5b)**. In all, this validation set included low, medium, and high efficiency pegRNAs spanning a range of 0 to 86% correct editing percentages, as measured by their respective sensor sites **(Fig. 5c)**. We transduced A549-PEmax cells with lentiviruses encoding individual pegRNAs and allowed editing to occur for 7–10 days, based on the kinetics of editing we observed previously **(Fig. 1f)**. We then mixed each individual population of sequence-verified isogenic A549-PEmax-pegRNA cells with parental *TP53* wild type A549-PEmax cells and performed longitudinal fluorescence competition assays in the presence or absence of Nutlin-3 **(Fig. 5d, Extended Data Fig. 7)**. We used the RFP fluorophore linked to each pegRNA vector to track the relative fitness of pegRNA cells (RFP+) compared to parental cells (RFP-) **(Fig. 5d).** These competition assays proceeded for two weeks, with flow cytometry readings taken every 7 days for each replicate **(Fig. 5d, Extended Data Fig. 8)**. We then calculated the difference in the RFP+ cell fraction (ΔRFP+ %) for each pegRNA between the final and initial time-point in both conditions **(Fig. 5e).** Consistent with our screening results, a significant fraction of pegRNAs showed enrichment in the presence of Nutlin-3, often reaching complete saturation **(Fig. 5e)**. Overall, there was strong concordance between the enrichment in Nutlin-3-treated cells observed in the screen and in these competition assays, supporting the reproducibility of the screening results **(Fig. 5f).** Importantly, we observed a significantly strong enrichment of cells expressing pegRNAs designed to engineer variants in the oligomerization domain, including A347P **(Fig. 5d, 5f).**

**Fig. 5.**
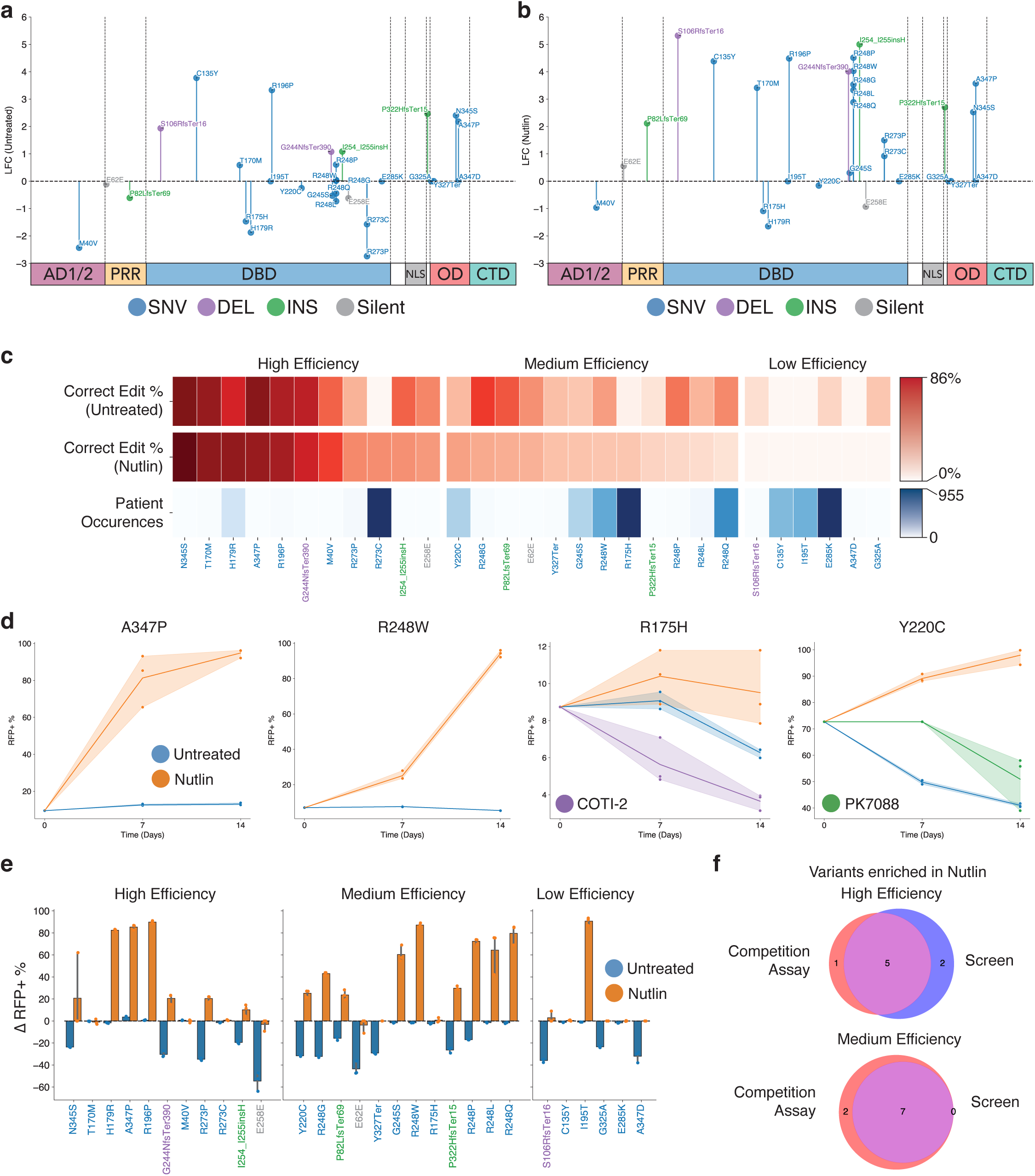
Functional validation of pathogenic *TP53* variants identified with prime editing screening. **a,** LFC of 29 pegRNAs selected for validation at Day 34 in the untreated condition. Variants with insufficient control counts (I195T, E285K, G325A, Y327Ter, A347D) represented with LFC = 0. **b,** LFC of 29 pegRNAs selected for validation at Day 34 in the Nutlin-treated condition. Variants with insufficient control counts (I195T, E285K, G325A, Y327Ter, A347D) represented with LFC = 0. **c,** Visualization of correct editing percentages and number of patient occurrences (MSK-IMPACT) for the pegRNAs selected for validation. **d,** Selected competition assays show the change in the RFP positive cell fraction in the presence or absence of various selection agents. **e,** The change in the RFP positive cell fraction (ΔRFP+ %) at the initial versus final time-point for each variant in the presence or absence of Nutlin-3. **f,** Venn diagram visualizing the agreement between enrichment in the presence of Nutlin in the competition assays (ΔRFP+ % 2: 5) and the original screen (LFC 2: 1) for high and medium efficiency pegRNA validation sets.

The above results indicate that cells harboring a number of pegRNAs designed to engineer diverse types of *TP53* variants have an increased fitness. However, these results do not rule out the possibility that these pegRNAs confer increased fitness through indel generation, rather than through their encoded edits. To assess this, we performed targeted Sanger sequencing and NGS across each of the target loci in A549-PEmax-pegRNA cells. Consistent with our screening data, sensor measurements, and competition assay results, each edited cell line exhibited specified on-target prime editing, in some instances reaching apparent homozygosity **(Extended Data Fig. 7)**.

Another potential application of our approach is to combine high-throughput prime editing with drug treatments to identify variant-drug interactions that could be exploited to develop allele-specific therapies. This is particularly relevant today because recent advances in rational drug design have shown that small molecules targeting specific mutant proteins (including those produced by oncogenic point mutant *KRAS* alleles) can have significant therapeutic potential^50^. To test if our approach could be used to identify variant-specific therapeutic sensitivities, we tested two mutant-p53-specific therapeutic agents, COTI-2 and PK7088, in cells transduced with lentiviruses encoding R175H- and Y220C-targeting pegRNAs, respectively^51–53^. Both of these treated populations showed depletion in the RFP+ cell fraction **(Fig. 5d).** In particular, the COTI-2 treatment arm showed significant depletion of R175H-pegRNA cells **(Fig. 5d).** These results demonstrate that prime editing sensor screens could be used to systematically identify variant-specific vulnerabilities to diverse therapies, augmenting cDNA-based approaches for performing similar screens^54^.

### Comparative analysis of prime editing and cDNA screening datasets reveals new pathogenic variants in the p53 oligomerization domain

High-throughput functional genomics approaches have been used previously to investigate *TP53* mutations. For instance, Giacomelli et al. performed deep mutational scanning of *TP53* variants using exogenous overexpression of mutant *TP53* cDNAs in A549 cells in the presence or absence of Nutlin-3, concluding that most *TP53* mutations likely arise as a consequence of endogenous mutational processes that select for dominant negative and LOF activity^6^. A follow up study integrated this method with HDR-based modeling of six *TP53* hotspot mutations in human leukemia cells and concluded that missense mutations in the *TP53* DBD act solely through dominant negative activity^2^. More recently, Ursu et al. employed a modified version of Perturb-seq to interrogate the transcriptional effects of 200 mutant *TP53* cDNAs in A549 cells by single cell RNA-sequencing, also concluding that most of these disrupt p53 activity through LOF and dominant negative effects^16^. In contrast, another study used a similar approach in H1299 and HCT116 cell lines to interrogate variants in the *TP53* DBD through parallel *in vitro* and *in vivo* experiments, concluding that certain hotspot mutations confer a higher proliferative advantage *in vivo*, likely through GOF mechanisms^7^. A number of studies in both mice and humans have also demonstrated that certain *TP53* variants, including hotspot mutations at residues R175, R248, and R273, can produce phenotypes consistent with neomorphic/GOF activities^55^. These include promoting aberrant self-renewal of hematopoietic stem cells^56^, sustaining tumor growth^57,58^, and promoting metastatic dissemination^59–62^, among others^55^. As such, there is much controversy in the field regarding the precise cellular and molecular activities of cancer-associated *TP53* mutations — the most common genetic lesions observed across all types of cancer.

We hypothesized that cDNA screening approaches are biased in favor of detecting dominant negative activities due to their reliance on supraphysiological overexpression of mutant proteins. This is particularly relevant for studying the active p53 transcription factor, which is a tetrameric protein composed of a dimer of dimers^18^. As such, we postulated that mutant overexpression studies in wild type *TP53* cells, including A549, may be incapable of detecting mutant allele-specific activities and phenotypes that may be sensitive to gene dosage and protein stoichiometry. To test this hypothesis, we reanalyzed our data to perform comparative bioinformatic analyses with the dataset generated by Giacomelli et al.^6^, as their experiments were also carried out in wild type *TP53* A549 cells treated with Nutlin-3. First, we plotted the Z-scores of SNV-generating pegRNAs (≥10% editing) against the Z-scores of the corresponding variants expressed from cDNAs **(Fig. 6a).** Supporting our hypothesis, variants in the OD of p53 tended to deplete (i.e., Z-score<0) when expressed from cDNAs, but often enriched significantly when expressed from the endogenous locus **(Fig. 6a, 6b)**. To investigate this difference further, we calculated the difference in Z-scores (ΔZ-score) between each pegRNA–cDNA pair by subtracting the prime editing Z-score from the cDNA Z-score. This analysis revealed a significantly higher ΔZ-score for endogenous variants in the OD relative to other domains of p53 **(Fig. 6c, 6d)**, a pattern that consistently held at multiple thresholds of pegRNA activity, and even when we restricted our analysis solely to the most efficient pegRNA for each variant **(Fig. 6e, 6f).** Some of the most impactful variants in the p53 OD are frequently observed in individuals with Li-Fraumeni syndrome, who carry germline *TP53* variants that predispose them to cancer. Importantly, recent studies by the Prives and Lozano laboratories have shown that p53 proteins harboring A347D mutations (in the OD) form stable dimers instead of tetramers, and that these dimeric p53 proteins exhibit neomorphic activities^63,64^. Lastly, visualizing the residue-averaged ΔZ-scores on the structure of the p53 OD^65^ further highlights the extensive differences in the behavior of endogenous variants as compared to exogenous (cDNA) variants in this domain **(Fig. 6g).** Collectively, our observations highlight gene dosage, protein stoichiometry, and protein-protein interaction domains as important variables that must be taken into account when studying the enormous diversity of mutant alleles observed in human cancer. Failing to take these considerations into account might lead to the misclassification of *bona fide* pathogenic variants, including those identified in patients with hereditary cancer predisposition syndromes like Li-Fraumeni.

**Fig. 6.**
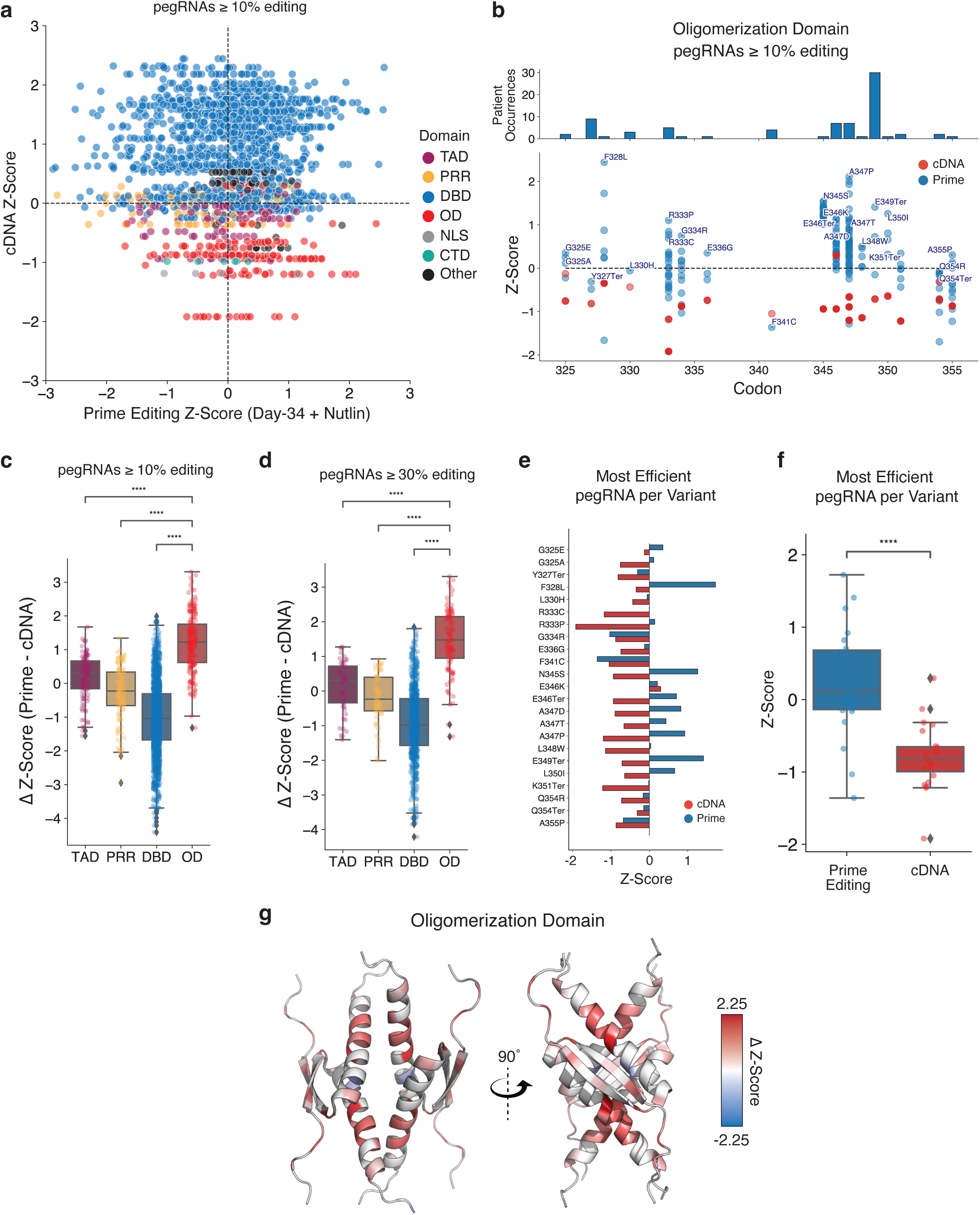
Comparative analysis of prime editing and cDNA screening datasets of *TP53* variants reveals novel pathogenic variants in the oligomerization domain. **a,** Scatter-plot of prime editing Z-score for pegRNAs 2: 10% editing at Day 34 in the Nutlin-treated condition, and the corresponding cDNA variant Z-score in the p53-WT background in the presence of Nutlin, colored by p53 domain. **b,** cDNA (red) and prime editing (blue) Z-scores for pegRNAs/variants located in the oligomerization domain. The pegRNA with the highest Z-score is labeled. **c, d,** The difference in Z-scores between prime editing and cDNA screens (ΔZ-score) for pegRNAs 2: 10% and 2: 30% editing, separated by p53 domain. Statistics shown for t-test of independent samples with Bonferroni correction. * = p-value ::5 .05, ** = p-value ::5 .01, *** = p-value ::5 .001, **** = p-value ::5 .0001, ns = not significant (p-value > .05). **e,** Z-scores for variants in the cDNA (red) and prime editing (blue) screens, considering only the most efficient pegRNA for each variant with an editing efficiency 2: 10%. **f,** Box-plot of the z-scores for variants in the cDNA (red) and prime editing (blue) screens, considering only the most efficient pegRNA for each variant. Statistics shown for t-test of independent samples with Bonferroni correction; **** = p-value ::5 .0001. **g,** Visualization of the residue-averaged ΔZ-scores (pegRNAs 2: 10% editing) on an NMR-structure of p53 oligomerization domain (PDB: 1OLG).

## Discussion

The overwhelming majority of genetic variants associated with various human diseases, including cancer, remain uncharacterized^40^. The development of CRISPR-based precision genome editing tools, including base and prime editing^9,10^, has opened the door to rigorous experimental interrogation of disease-associated variants with single base-pair resolution at unprecedented scales. However, these approaches, particularly prime editing, remain limited by the variance in editing efficiency among different pegRNAs.

Building on our previous work, we developed a new prime editing sensor-based framework for engineering and screening variants that overcomes these limitations. Using this sensor-based approach, in which each pegRNA is coupled to an artificial version of its endogenous target site, we simultaneously identify enriched pegRNAs while empirically quantifying their editing efficiencies. This novel approach allows for the characterization of variants at multiple target sites while correcting for the potentially confounding differences in editing activity among distinct pegRNAs. To facilitate the creation of similar libraries by other researchers, we also built PEGG, a new computational tool for creating prime editing sensor libraries (<u>pegg.readthedocs.io</u>).

As a prototype for the prime editing sensor framework, we generated a library of pegRNAs targeting over a thousand cancer-associated variants in *TP53*. We reasoned that p53 would be the most salient prototype for testing the efficacy of our prime editing sensor screening platform because of its central role in cancer development and progression, the various studies that have performed deep mutational scanning of p53 whose data can provide direct sources of comparison, and the ongoing controversy in the field as to whether most (if not all) *TP53* mutations observed in cancer patients are functionally redundant.

With our sensor-based approach, we were able to systematically search the pegRNA design space and identify high efficiency pegRNAs, allowing us to install over half of the targeted *TP53* variants. Though not the focus of the present study, this dataset also allowed us to recapitulate many of the previous findings about the factors affecting pegRNA efficiency.

Analysis of the screening data revealed a wide range in the fitness of *TP53* variants, challenging the idea that most p53 variants, particularly those in the DBD, are functionally redundant. We identified strongly enriching pegRNAs, including pegRNAs that generated “hotspot” mutations that are commonly observed in patients, particularly those at DNA-contacting residues. However, the majority of the strongly enriching pegRNAs encoded variants that are not located at mutational hotspots; instead, a number of these mutations are less frequently observed in patients and remain poorly understood, despite the fact that they affect thousands of patients globally every year. These include variants located within the OD of p53, which were functionally validated with follow-up experiments and some of which were recently shown to be *bona fide* pathogenic variants in humans with Li-Fraumeni Syndrome^63,64^.

Comparison between our screening data and a previous study that also screened *TP53* variants in A549 cells and in the presence of Nutlin-3 but instead used cDNA-based overexpression libraries revealed a statistically significant enrichment of endogenous — but not overexpressed — variants located in the p53 OD. This comparison highlights the potential limitations of cDNA-based screening approaches, particularly when studying variants at the sites of protein-protein interactions, underscoring the need to study variants in their native context to access their true biology. Together, our data suggest that stoichiometric imbalances produced by cDNA overexpression could lead to the mis-classification of genetic variants as non-causal or otherwise benign. Our findings thus offer a cautionary note when using exogenous overexpression systems to interpret pathogenic alleles, and highlight the importance of using strategies like the one described in this work to investigate variants of interest in their native genomic contexts whenever possible.

More broadly, our study provides a conceptual blueprint and a modular set of experimental and computational tools that can be applied to evaluate diverse types of genetic variants in their native endogenous genomic contexts with high-throughput prime editing. Future efforts could incorporate improved prime editors and pegRNAs, as well as higher-efficiency prime editing systems such as PE3 and PE4^10,21^. These studies could also be performed *in vivo*, for example by delivering compact libraries of MMR-evasive pegRNAs into mice expressing prime editors^66^ or by co-delivery of smaller prime editors, such as PE6a^23^. Taken together, we envision that our approach will expand our understanding of pathogenic gene variants and help match patients suffering from genetic diseases with effective therapies.

## Methods

### Experimental Materials & Methods

#### Plasmids and pegRNA cloning

All plasmids were generated using Gibson Assembly strategies^67^ using NEBuilder^®^ HiFi DNA Assembly Master Mix (NEB #E2621) following the manufacturer’s protocol. All new plasmids, along with detailed maps and sequences, will be made available through Addgene. The PEmax coding sequence in Lenti-EFS-PEmax-P2A-Puro was obtained from pCMV-PEmax (Addgene, catalog no. 174820)^21^. The lentiviral plasmid used to clone and express prime editing sensor libraries was assembled by transferring the U6-sgRNA-EFS-Blast-P2A-TurboRFP cassette from pUSEBR^14^ into the higher titer pLV backbone^68^. Lenti-PEAR-mCherry, a modified all-in-one lentiviral version of the PEAR^43^ reporter, was also cloned using Gibson Assembly and used to test the editing activity of A549-PEmax cells. The Lenti-UPEmS-tevo plasmid is a modified version of the UPEmS vector^66^ that contains the tevopreQ1 motif^22^. This plasmid was used to assemble pegRNAs via Golden Gate Assembly^66^ for follow-up pegRNA validation experiments.

#### Virus Production

Lentiviruses were produced by co-transfection of HEK293T cells with the relevant lentiviral transfer vector and packaging vectors psPAX2 (Addgene, catalog no. 12260) and pMD2.G (Addgene, catalog no. 12259) using Lipofectamine 2000 (Invitrogen, catalog no. 11668030). Viral supernatants were collected at 48- and 72-hours post-transfection and stored at -80°C.

#### Drug treatments

Nutlin-3 (Selleck Chemicals, S1061) was dissolved in DMSO at a stock concentration of 10 mM and used at a final concentration of 10 μM. PK7088 (Aobious #AOB4255) was diluted to a final concentration of 200 µM from a stock concentration of 10 mM. COTI-2 (MedChemExpress #HY-19896) was dissolved in DMSO to a stock concentration of 10 mM and added to a final concentration of 1 µM.

#### Flow Cytometric Analyses

Fluorescence-based measurements for the validation of prime editing activity with PEAR and for competition assays were performed using the BD FACSCelesta Cell Analyzer in tube or plate reader format. Downstream analysis was performed using FlowJo to identify single cells and quantify fluorescence.

#### Generation of A549-PE Max cell lines

To generate A549 cells stably expressing PEmax, we transduced a 15-cm plate with 2.5 million cells with freshly harvested EFS-PEmax-P2A-Puromycin lentivirus, and selected cells with 10 µg/mL of puromycin at 72-hours post-transduction. To assess the prime editing efficiency of these cells, we transduced 250K A549-PEmax cells in triplicate in 6-well plates with Lenti-PEAR-mCherry, a modified, all-in-one lentiviral PEAR construct, where GFP is turned on in the event of successful prime editing. We noticed that prime editing activity was not sufficiently high in these cells based on the PEAR reporter activity. We then re-transduced these cells with 2 successive rounds of freshly harvested EFS-PEmax-P2A-Puromycin lentivirus. Repeating the PEAR reporter assay revealed a substantial increase in prime editing activity **(Extended Data Fig. 2a)**. This A549-PEmax “v2” cell line was used throughout the present study.

#### Cloning of p53-sensor Library

The oligonucleotide library was ordered from Twist Biosciences. The lyophilized library was resuspended in 100 µL of TE Buffer (pH 8.0) and diluted to create 1 ng/µL stocks. We performed n=32 PCR reactions with NEBNext High-Fidelity 2X PCR Master Mix (#M0541S) to amplify the library with the following primers at a low cycle count: (1) F: 5′-CATAGCGTACACGTCTCACACCG, (2) R: 5′-GTGCCGTTGACGACCGGATCTAGAATTC. These PCR reactions were pooled and PCR purified using the Qiagen PCR purification kit following the manufacturer’s protocols, with 10 µL of 3 M NaOAc pH 5.2 added for every 5 volumes of PB used per 1 volume of PCR reaction. The library was digested with Esp3I (NEB) and EcoRI-HF (NEB), pooled and PCR purified. Subsequently, n=16 ligations were performed using 300 ng of digested and dephosphorylated Trono-BR backbone and 3 ng of digested insert with high concentration T4 DNA Ligase (NEB #M0202M). The ligation reactions were precipitated using QuantaBio 5PRIME Phase Lock Gel tubes before being resuspended in 3 µL of EB Buffer per 4 precipitated reactions. These precipitated ligation reactions were electroporated into Lucigen Endura ElectroCompetent Cells (#60242-2) before being plated on LB-Carbenicillin plates and incubated at 37°C for 16 hours. Library representation was assessed at this step via serial dilution plating, displaying a representation on the order of 400x. We also picked 30 random colonies from these serial dilution plates to assess the fidelity of library cloning and to test a random set of pegRNAs. We scraped the plates and collected the bacteria in 250 mL of LB-Ampicillin per 4 plates, before incubating for 2 hours at 37°C, spinning the bacteria, and proceeding to perform a Qiagen Maxiprep, following the manufacturer’s protocol. Lentivirus was generated via the aforementioned protocol, and viral titer was determined through serial dilutions of virus, transductions in 12-well plates with 1 million A549-PEmax cells, and measurement of the RFP-positive cell fraction at 72-hours post-transduction. For extended protocol information, see **Supplemental Protocol 1.**

#### Screening protocol

For each replicate, 110 million A549-PEmax cells were combined with an appropriate amount of p53-sensor virus to achieve an MOI < 1. To this mix, puromycin was added to a final concentration of 10 µg/mL and polybrene transfection reagent (Sigma-Aldrich, catalog no. TR-1003) was added to a final concentration of 8 µg/mL in F-12K (Gibco, catalog no. 21127030) media supplemented with 10% FBS and 1X Penicillin-Streptomycin (ThermoFisher). This mix was plated into 9 total 12-well plates per replicate. At 24-hours post-transduction, each 12-well plate was expanded to 15-cm plate, with media supplemented with 10 µg/mL puromycin and 10 µg/mL blasticidin S. These puromycin and blasticidin S concentrations were maintained throughout the screen. At 72-hours post-transduction, each 15-cm plate was expanded to 2 15-cm plates. At 96-hours post-transduction, each replicate was replated at ≥1000X representation (≥29 million cells) for the untreated arm and the Nutlin-3 treatment arm. For the Nutlin-3 treatment arm, Nutlin-3 was added to a final concentration of 10 µM. At this time-point, a cell pellet was taken for gDNA extraction. All cell pellets included ≥29 million cells (1000X representation) and were stored at –80°C. Subsequently, every 3 days, the cells were split, and replated at 1000X representation. At each time-point, a cell pellet was taken if there were a sufficient number of cells to allow for 1000X representation. This process was repeated until the screen was terminated at Day 34 post-transduction.

#### Genomic DNA extraction

Genomic DNA from the Day 4, 16, 25, and 34 time-points was extracted using the Qiagen Genomic Tip/500 G following the manufacturer’s protocol. Genomic DNA was resuspended in 200 µL of TE Buffer, pH 8.0. Concentrations were measured using a NanoDrop 2000 (ThermoFisher) and were normalized to 1 ug/uL.

#### NGS Sample Preparation

We performed n=30 PCR1 reactions per sample using Q5 High Fidelity 2X Master Mix (NEB #M0429S) with 10 ug of genomic DNA to maintain ≥1000X representation. Up to 4 PCR reactions were pooled and PCR purified using the Qiagen PCR purification kit following the manufacturer’s protocols. These reactions were then gel purified (Qiagen Gel Extraction Kit), pooled and measured using the NanoDrop 2000 (ThermoFisher). We performed n=4 PCR2 reactions per sample using 10 ng of PCR1 as a template in each reaction. We PCR purified these samples, and then gel purified these samples and eluted in 30 µL of EB Buffer. These samples were then submitted for sequencing. The PCR1 and PCR2 strategies are diagrammed in **Extended Data Fig. 9a**. All primers are deposited in **Supplemental Table 2.**

#### Next Generation Sequencing (NGS)

We performed Amplicon-EZ sequencing (Azenta) for analysis of the correlation between sensor and endogenous editing **(Fig. 1d).** For the NGS of the p53-sensor library, we used the NovaSeq S1 200 sequencing system (NovaSeq 6000) with a custom sequencing primer set to amplify the protospacer, 3′ extension, sensor sequence, and sample barcode in separate reads. All other NGS data was generated using the Singular G4 Sequencer (2×150 paired-end) with stock primers. All sequencing primers are deposited in **Supplemental Table 2.** The custom sequencing approach for the NovaSeq 6000 is diagrammed in **Extended Data Fig. 9b**.

#### Golden Gate Assembly of UPEmS pegRNAs for follow-up validation

For follow-up validation of pegRNAs, individual pegRNAs were cloned via Golden Gate Assembly into the Lenti-UPEmS-tevo backbone (generated by the present study). Golden Gate Assembly was performed with annealed spacer oligonucleotides, annealed and phosphorylated scaffold oligonucleotides, and annealed 3′ extension oligonucleotides using NEB BsmBI Golden Gate Enzyme Mix, before being transformed, mini prepped (Qiagen), and validated via whole-plasmid sequencing (Primordium). For full protocol details, see **Supplemental Protocol 2**.

### Competition assays

To generate variant p53 lines, we seeded 100K A549-PEmax cells in 6-well plates and added UPEmS lentivirus corresponding to each variant. To achieve saturation editing, we waited 7–10 days, expanding the cells to a 10-cm plate when they reached confluence. At this point, we mixed 250K variant (RFP+) cells with 750K un-transduced A549-PEmax cells, and plated 50K cells in triplicate in 6-well plates. For drug treated conditions (Nutlin-3, COTI-2, PK7088), the compound was added to the appropriate concentration. Remaining cells were used for flow analysis and to generate a t=0 cell pellet. At day 7 and day 14, we assessed the RFP positive fraction of the cells via flow cytometry, and at the final time-point took a cell pellet for gDNA extraction.

### Analytic/Computational Methods

#### Selection of *TP53* Variants and prime editing sensor library generation with PEGG

To select a cohort of *TP53* variants for generating a prime editing sensor library, we used the MSK-IMPACT database^20^. We chose all SNVs observed in patients, as well as a collection of observed and random indels to increase the diversity of edits **(Extended Data Fig. 1b)**. In addition, PEGG automatically generated 95 neutral/silent variants that tiled the *TP53* locus to act as internal controls in the screen.

PEGG generated a maximum of 30 ranked pegRNA designs per variant with RTT lengths of 10, 15, 20, 25, and 30 nt, and PBS lengths of 10, 13, and 15 nt coupled to “NGG” protospacers. A “G” was appended to the start of each 20 nt protospacer to improve U6 promoter-mediated transcription. After PEGG generated these pegRNA designs, we further filtered the library to exclude pegRNAs containing polyT termination sequences (≥ 4 consecutive Ts), EcoRI and Esp3I sites, and protospacers with an MIT specificity score less than 50. In addition, each pegRNA oligo included a matched, 60 nt sensor locus that was automatically generated by PEGG and used to link each pegRNA to its editing outcome.

For full details of generating a prime editing sensor library using PEGG, visit pegg.readthedocs.io.

#### Analysis of the p53-sensor screen

The p53-sensor sequencing results were demultiplexed into separate fastq files based on the sample barcode. Next, using a custom analysis script, we filtered reads with an average Phred quality score below 30, and identified pegRNAs based on the protospacer and 3′ extension sequences. Sequences with no matching protospacer or 3′ extension were discarded, and sequences with mismatched protospacer and 3′ extension sequences were discarded and classified as recombination events. Sequences with matching protospacer and 3′ extension sequences were used to generate pegRNA counts tables that were subsequently used for MAGeCK analysis of pegRNA enrichment/depletion.

To classify editing outcomes at the sensor locus, we first determined whether recombination had occurred to decouple the pegRNA from its matched target sequence. To do so, we used the first and last 5-nt of the sensor sequence as a barcode to detect recombination. Sensor reads with the first and last 5-nt of the read matching the appropriate pegRNA were classified as correct sensor reads, while those with the first and last 5-nt matching other pegRNAs were classified as recombination events and discarded. We noted that recombination between the pegRNA and sensor was observed at a significantly higher rate when the protospacer was in the same orientation as the sensor, prompting us to update PEGG to automatically place the sensor sequence in the reverse orientation to reduce recombination in future PE sensor libraries. **(Extended Data Fig. 2e, 2f).** Reads with the first and last 5-nt with no match were classified as potential indels and retained. For each sample, the sensor reads that were not recombined were demultiplexed into separate fastq files for each pegRNA. We then used Crispresso2^69^ to classify editing outcomes, excluding the first and last 5-nt of the sensor read from the quantification window. To determine the background subtracted correct editing %, we subtracted the correct editing % observed in the plasmid library from the correct editing % observed at a given time-point, although we note that for plasmids with at least 10 sensor reads, the median correct editing % was 0%, the average correct editing % was < 0.1%, and the max observed correct editing % was 8.7%.

For analysis of enrichment/depletion of pegRNAs, we used the MAGeCK algorithm^47^, with the Day 4 sample designated as the control time-point. We then filtered to exclude pegRNAs with a control count mean <10 reads to reduce spuriously enriching pegRNA hits. For direct comparison with the cDNA libraries, the LFC values produced by MAGeCK were transformed into Z-Scores using the standard Z-score formula including all pegRNAs under consideration.

#### Processing and Analysis of *TP53* NGS Sequencing

The sequencing files were automatically demultiplexed into separate fastq files based on the sample barcode (12 total samples). Next, we trimmed the sequences from 150 nucleotides to 100 nucleotides, to allow the sequences to be joined. The sequences were joined using the fastq-join algorithm with the default parameters enabled^70^. We then used custom analysis scripts to generate counts tables for all of the unique sequences, merge matching samples from different flow cells, and determine the HGVSp and HGVSc of each sequence.

To determine the LFC of each variant at these endogenous loci, we first filtered to exclude undesired variants (i.e. those not targeted by pegRNAs in the library) and created counts tables for Day 4, Day 34 (untreated) and Day 34 (Nutlin-treated) for each of the three amplicons. For each amplicon, we used MAGeCK to normalize read counts between samples and determine the LFC of each variant. We then filtered endogenous variants with a control count <10 reads to reduce spuriously enriching variants. These MAGeCK tables from the different amplicons were concatenated to perform downstream analysis, comparing the endogenous variants with the pegRNA–sensor sequencing results. In addition, we performed sequencing of the WT A549 PEmax cell line, which confirmed the WT status of the regions amplified.

## Data Availability

Raw sequencing data from the screen is deposited in the SRA under accession PRJNA1014453. All processed datasets, analysis scripts, as well as Jupyter notebooks and source data for generating each figure that appears in the paper, are available at the following GitHub repository: https://github.com/samgould2/p53-prime-editing-sensor. Further documentation and installation instructions for PEGG are available at <u>pegg.readthedocs.io</u>.

## Data Tables

- **Supplemental Table 1:** pegRNAs & variant information
- **Supplemental Table 2:** Primers
- **Supplemental Protocol 1:** Cloning of p53 PE sensor libraries
- **Supplemental Protocol 2:** Golden Gate Assembly of pegRNAs in Lenti-UPEmS-tevo backbone
- Other information and source data are provided in the <u>Github Repository</u>.

## Acknowledgements

We dedicate this work in memory of Jingzhi Zhu, a research computing specialist at the Koch Institute Integrated Genomics and Bioinformatics Core who worked tirelessly to ensure that our community had a robust computing infrastructure. We thank Nicolas Mathey-Andrews, Carol L. Prives, Michael T. Hemann, Jonathan S. Weissman, Luke W. Koblan, and Lukas Dow for excellent scientific discussions and overall support. S.I.G. was supported by T32GM136540 and the MIT School of Science Fellowship in Cancer Research. F.J.S.R. is an HHMI Hanna Gray Fellow and was supported by the V Foundation for Cancer Research (V2022-028), NCI Cancer Center Support Grant P30-CA1405, the Ludwig Center at MIT (2036636), Koch Institute Frontier Awards (2036648 and 2036642) and the MIT Research Support Committee (3189800). This work was also supported in part by the Koch Institute Support (core) Grant P30-CA14051 from the National Cancer Institute. We also thank the Koch Institute Swanson Biotechnology Center for technical support, especially the Flow Cytometry Core, the Barbara K. Ostrom (1978) Bioinformatics Facility, and the Genomics Facility. A.H. is a National Science Foundation (NSF) Graduate Research Fellow. A.H. and D.R.L. were supported by NIH U01AI142756, R35GM118062, RM1HG009490, and HHMI. This article is subject to HHMI’s Open Access to Publications policy. HHMI lab heads have previously granted a nonexclusive CC BY 4.0 license to the public and a sublicensable license to HHMI in their research articles. Pursuant to those licenses, the author-accepted manuscript of this article can be made freely available under a CC BY 4.0 license immediately upon publication.

## Author Contributions

S.I.G. and F.J.S.R. conceived the project and wrote the manuscript. S.I.G. performed all computational analyses and analyzed experimental data. S.I.G., A.N.W., K.D., and G.A.J. performed experiments. G.A.J. generated Lenti-PEAR-mCherry. V.K.N. generated Lenti-EFS-PEmax-P2A-Puro. S.S.L. assisted in designing sequencing strategy and performing next-generation sequencing. A.H. and D.R.L. provided conceptual and technical advice and comments on the manuscript. F.J.S.R. supervised the work and secured funding.

---

**Extended Data Fig. 1.**
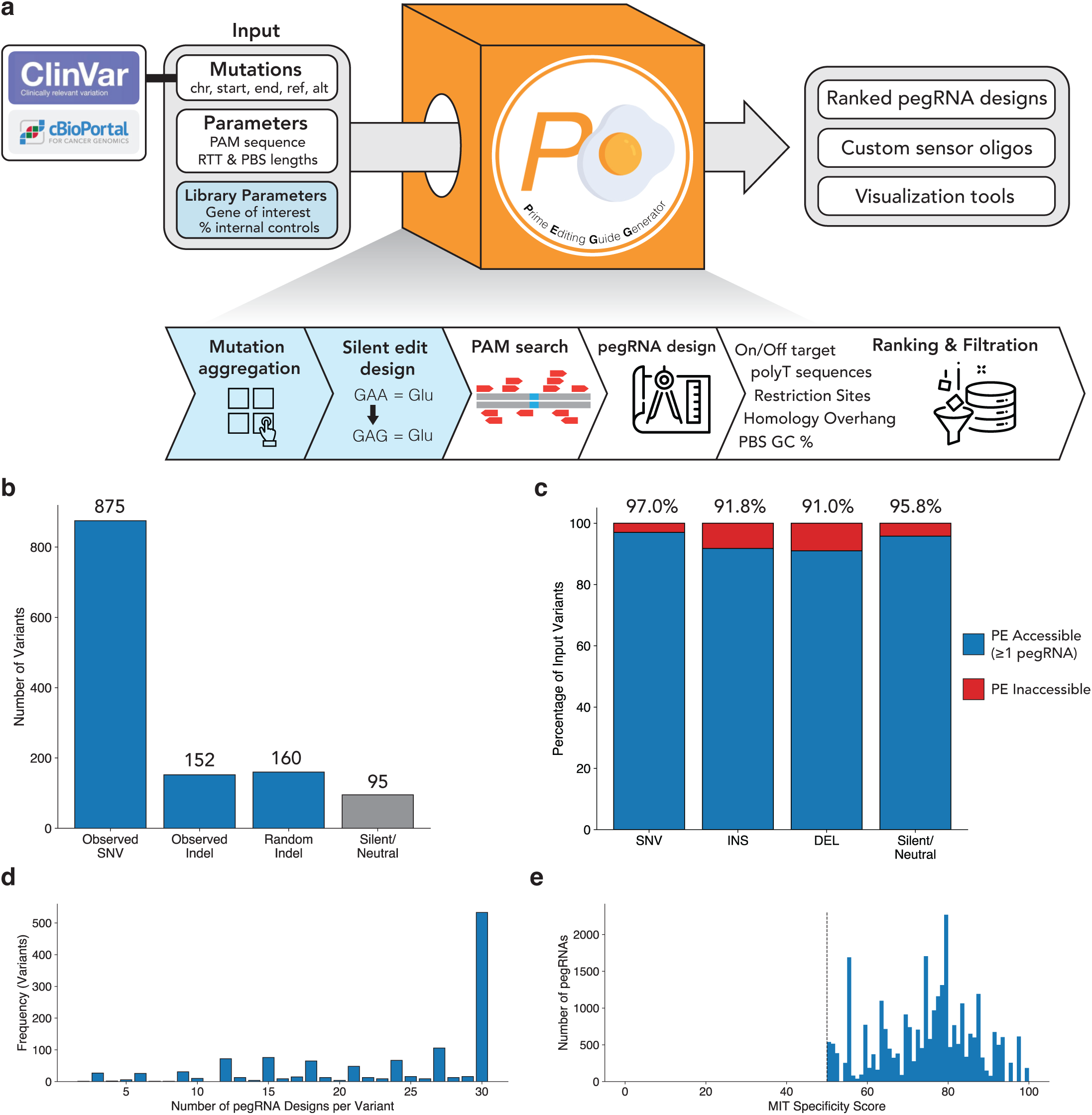
Generation of a TP53 prime editing sensor library with PEGG. **a,** Schematic of the PEGG pipeline. PEGG takes as input a list of mutations from the cBioPortal, ClinVar identifiers, or custom mutations sets, and produces user-defined pegRNA designs ranked by PEGG score, custom sensor oligos, and visualization tools. PEGG filters pegRNAs with polyT sequences as well as designs containing restriction sites used for cloning. With the library design feature, PEGG can also automate the aggregation of variants located in genes of interest, and automatically design a defined fraction of silent variant-generating pegRNAs. **b,** Breakdown of the TP53 variants input to PEGG for sensory library design. **c,** The fraction of input variants amenable to prime editing (i.e., able to generate ≥ 1 pegRNA), separated by variant type. **d,** Histogram of the number of pegRNA designs per variant. **e,** Histogram of the MIT specificity score of the protospacers for the pegRNAs included in the library. The library was filtered to exclude pegRNAs containing a protospacer with an MIT specificity score less than 50.

**Extended Data Fig. 2.**
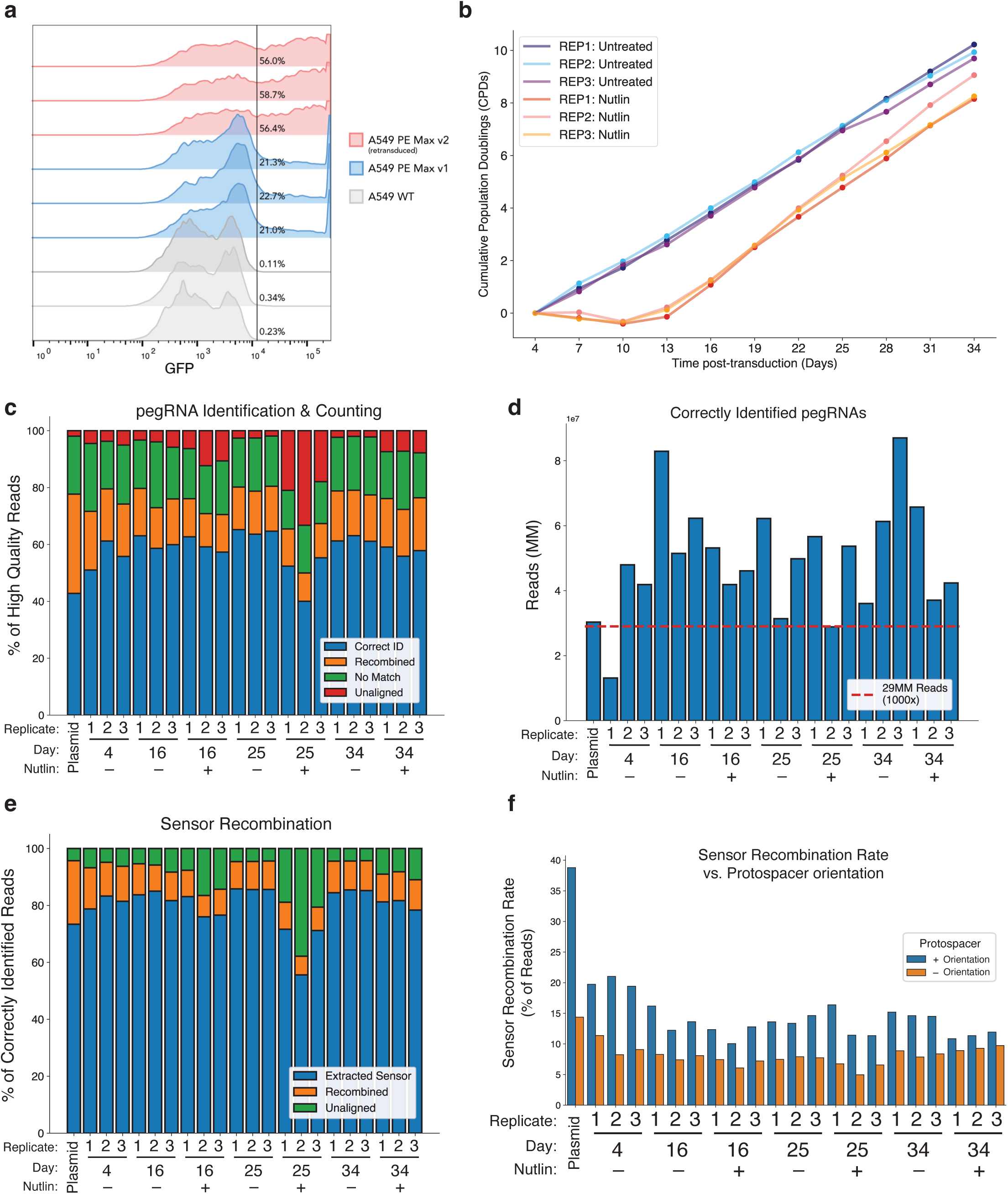
Optimization, screening, and deconvolution of the *TP53* prime editing sensor library. **a,** Assessment of the prime editing activity of A549-PEmax cell lines using a modified all-in-one PEAR reporter, where GFP is turned on in the event of successful prime editing. Three replicates each of A549 WT (gray), A549-PEmax v1 (blue), which underwent a single transduction with EFS-PEmax-P2A-Puro, and A549-PEmax v2 (pink), which underwent multiple rounds of transduction with EFS-PEmax-P2A-Puro, are shown, along with the quantification of GFP-positive cells. **b,** Cumulative population doublings during the course of the screen of each of the replicates in the untreated and Nutlin-treated conditions. **c,** Identification and counting of pegRNAs in each replicate and time-point from high quality reads (Q > 30). Correct ID = reads with matching protospacer and 3’ extension. Recombined = reads with mismatched protospacer and 3’ extension. No match = reads with no matching sequence for protospacer or 3’ extension. Unaligned = reads with no identifiable tevopreQ1 sequence. Plasmid = Plasmid Library. **d,** Count of correctly identified pegRNAs in each replicate and time-point. **e,** Extraction of the sensor locus from reads with correctly identified pegRNAs. Extracted sensor = sensor read matches pegRNA identification and is thus extracted and saved. Recombined = sensor read does not match pegRNA (discarded). Unaligned = no polyT-tevopreQ1 sequence found. **f,** Sensor recombination rate as a function of protospacer orientation. When the protospacer is on the positive strand (+) of the sensor (blue), the recombination rate increases compared to when the protospacer is on the negative strand (-) of the sensor.

**Extended Data Fig. 3.**
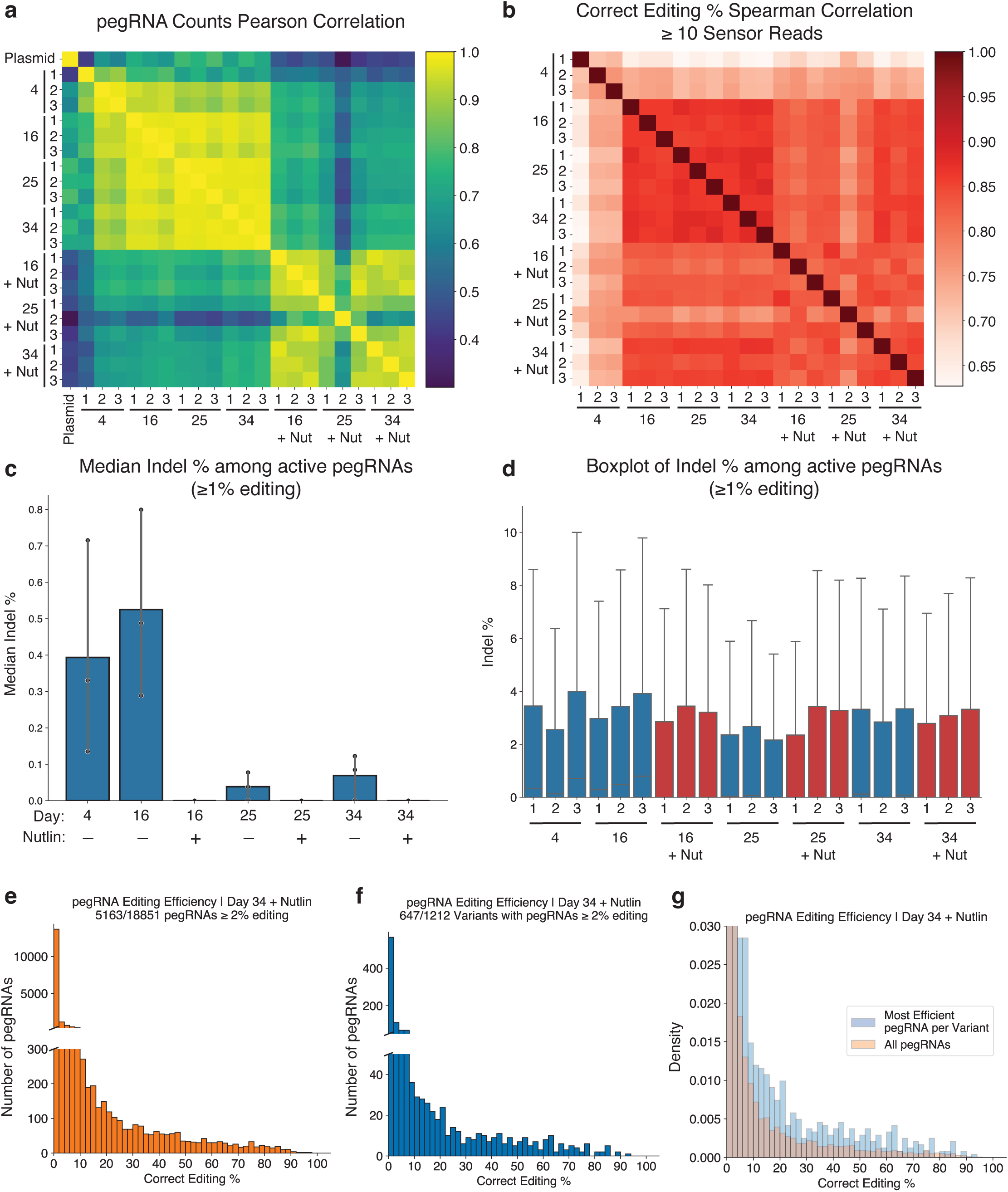
The *TP53* prime editing sensor screen is highly reproducible with low indel rates. **a,** Pearson correlation in raw pegRNA counts among each replicate and time-point. Plasmid = plasmid library. **b,** Spearman correlation in sensor correct editing percentage among each replicate and time-point for pegRNAs with at least 10 sensor reads. **c,** Median indel percentage among active pegRNAs (≥1% editing) for each time-point and condition. **d,** Boxplot of indel frequency among active pegRNAs (≥1% editing) for each replicate and time-point. Lower quartile = 0% for all replicates (not visible). **e,** Histogram of pegRNA editing efficiency in the Nutlin-treated Day 34 samples. **f,** Histogram of pegRNA editing efficiency of the most efficient pegRNA for each variant in the Nutlin-treated Day 34 samples. **g,** Comparative PDF of all pegRNAs and the most efficient pegRNA for each variant.

**Extended Data Fig. 4.**
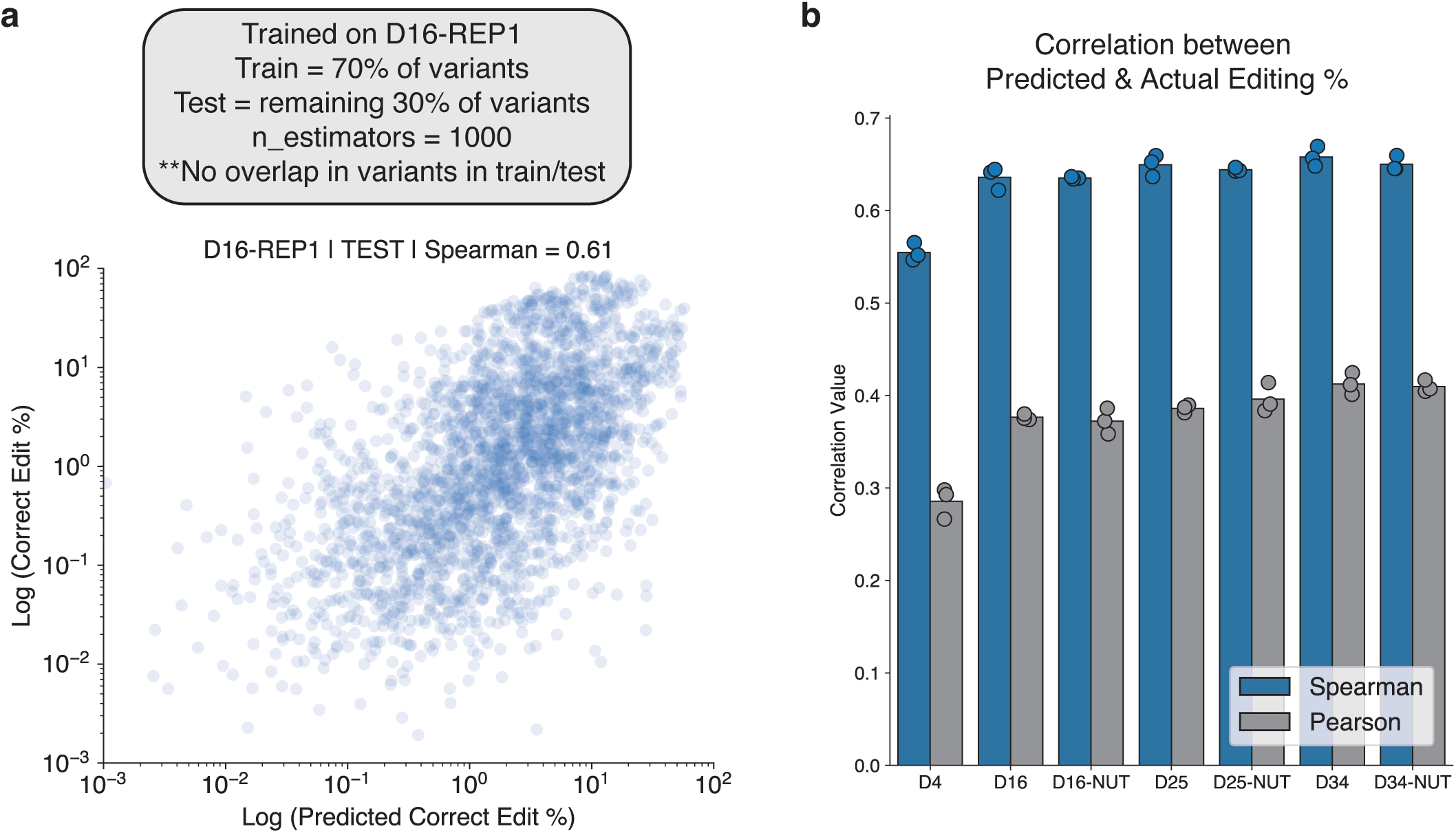
Training a random forest regressor to predict pegRNA efficiency. **a,** A random forest regressor was trained on a restricted set of pegRNA features using 70% of the variants in the untreated condition of Day 16 replicate 1. There was no overlap between the variants used for training and testing. The performance on the held-out test set is shown (spearman correlation = 0.61). **b,** Assessment of the performance of the random forest regressor in predicting editing activity at each time-point. Again, only variants in the test set are considered. Each dot represents a separate replicate. Spearman correlation between predicted and actual editing is shown in blue, pearson correlation in gray.

**Extended Data Fig. 5.**
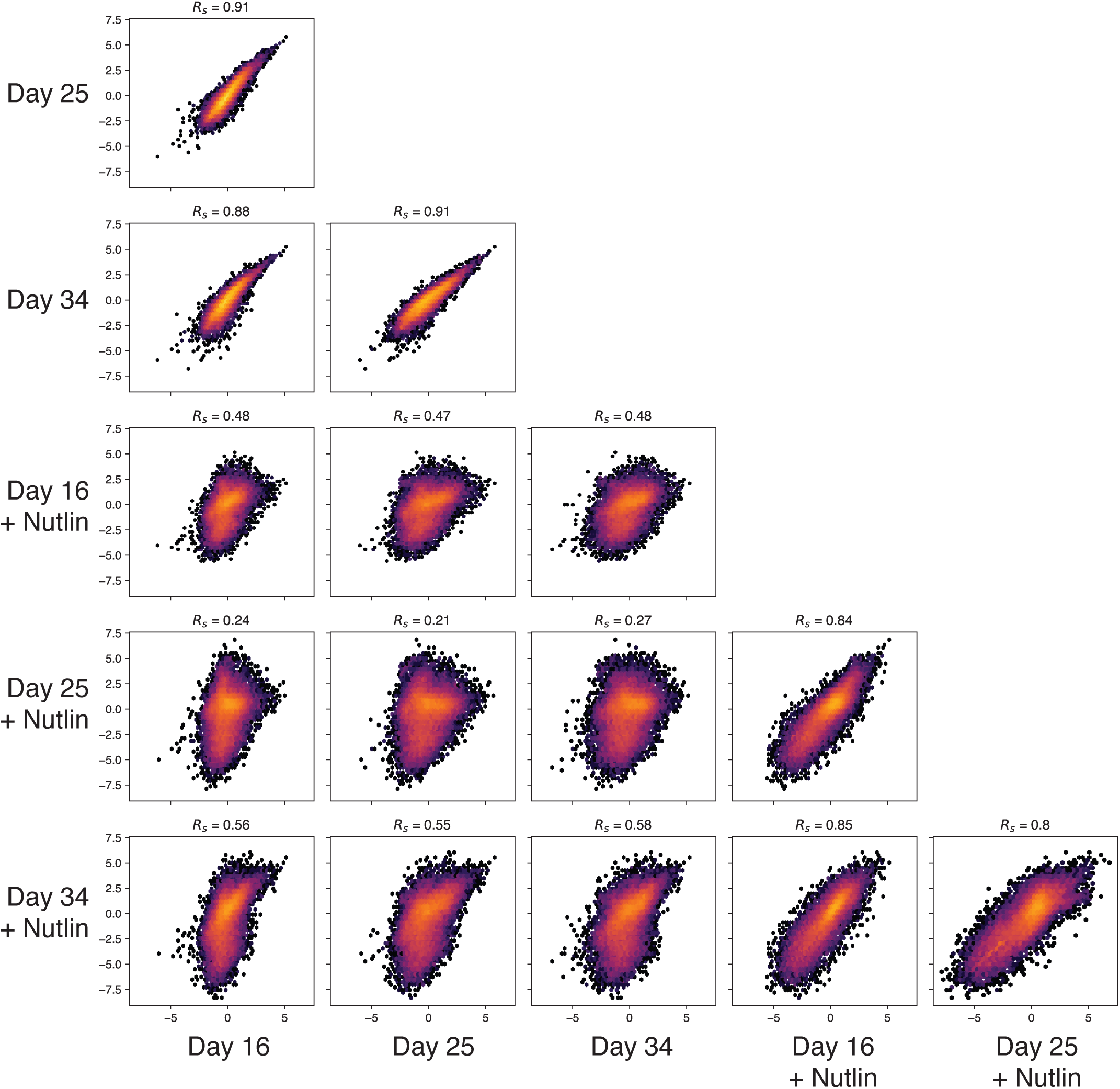
Correlation in pegRNA LFC among conditions and time-points. Each panel is a density plot of the LFC in pegRNAs at each time-point/condition (i.e. x-axis = LFC of the pegRNAs corresponding with that column’s sample, and y-axis = LFC of the pegRNAs corresponding with that row’s samples). Replicates were merged using MAGeCK to generate a single (median) LFC for each pegRNA at each time-point. Rs = spearman correlation.

**Extended Data Fig. 6.**
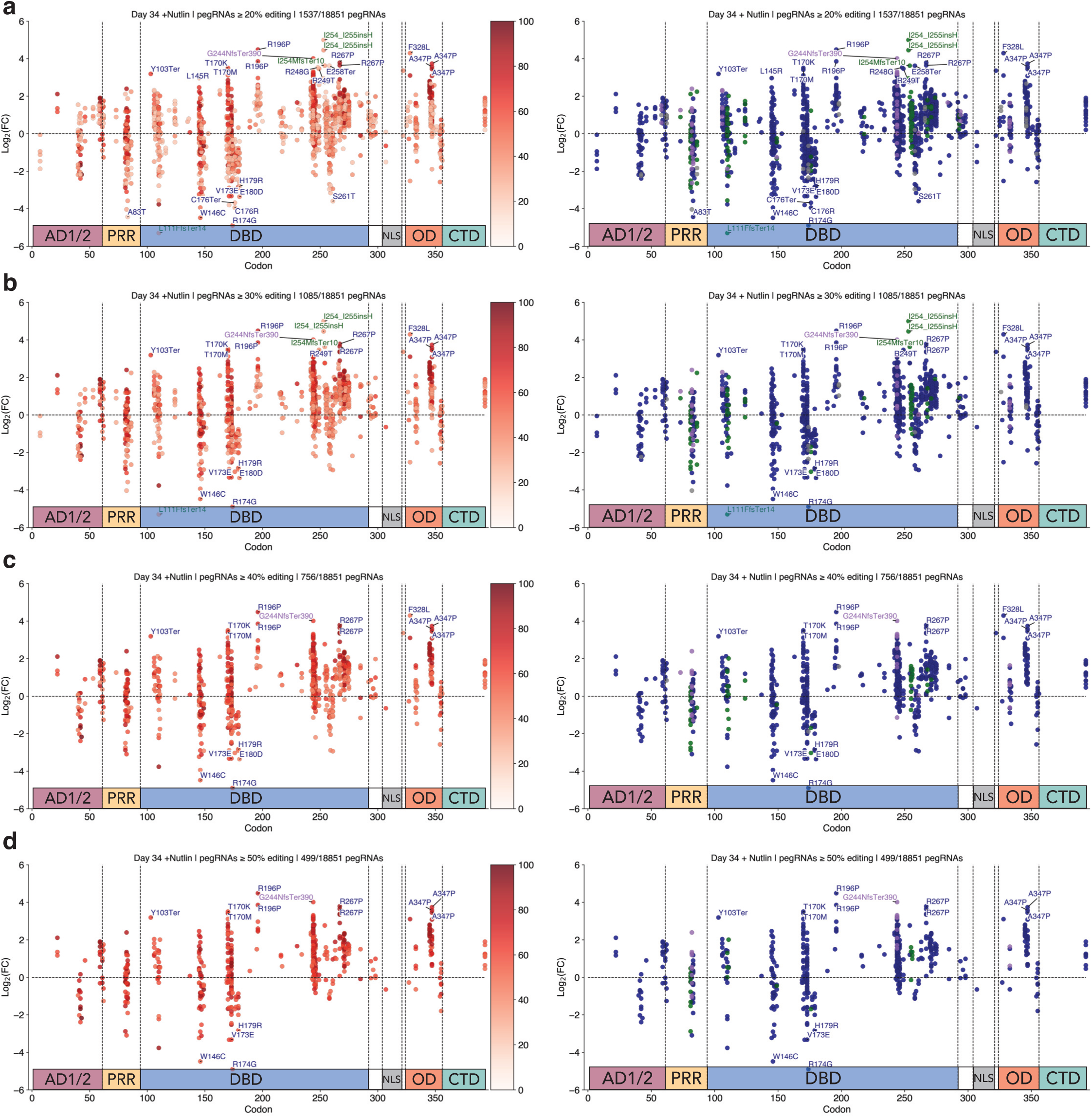
Filtration of screening data by sensor editing efficiency. The LFC of **a,** pegRNAs ≥ 20% editing **b,** pegRNAs ≥ 30% editing **c,** pegRNAs ≥ 40% editing **d,** pegRNAs ≥ 50% editing, with at least 10 sensor reads at Day 34 relative to Day 4 in the Nutlin-treated condition, with pegRNAs colored by editing efficiency (left) and colored by variant type (right). Selected enriching pegRNAs with FDR < .05 labeled and depleting pegRNAs with FDR < .05 labeled. Blue = SNV, Green = INS, Purple = DEL, Gray = Silent.

**Extended Data Fig. 7.**
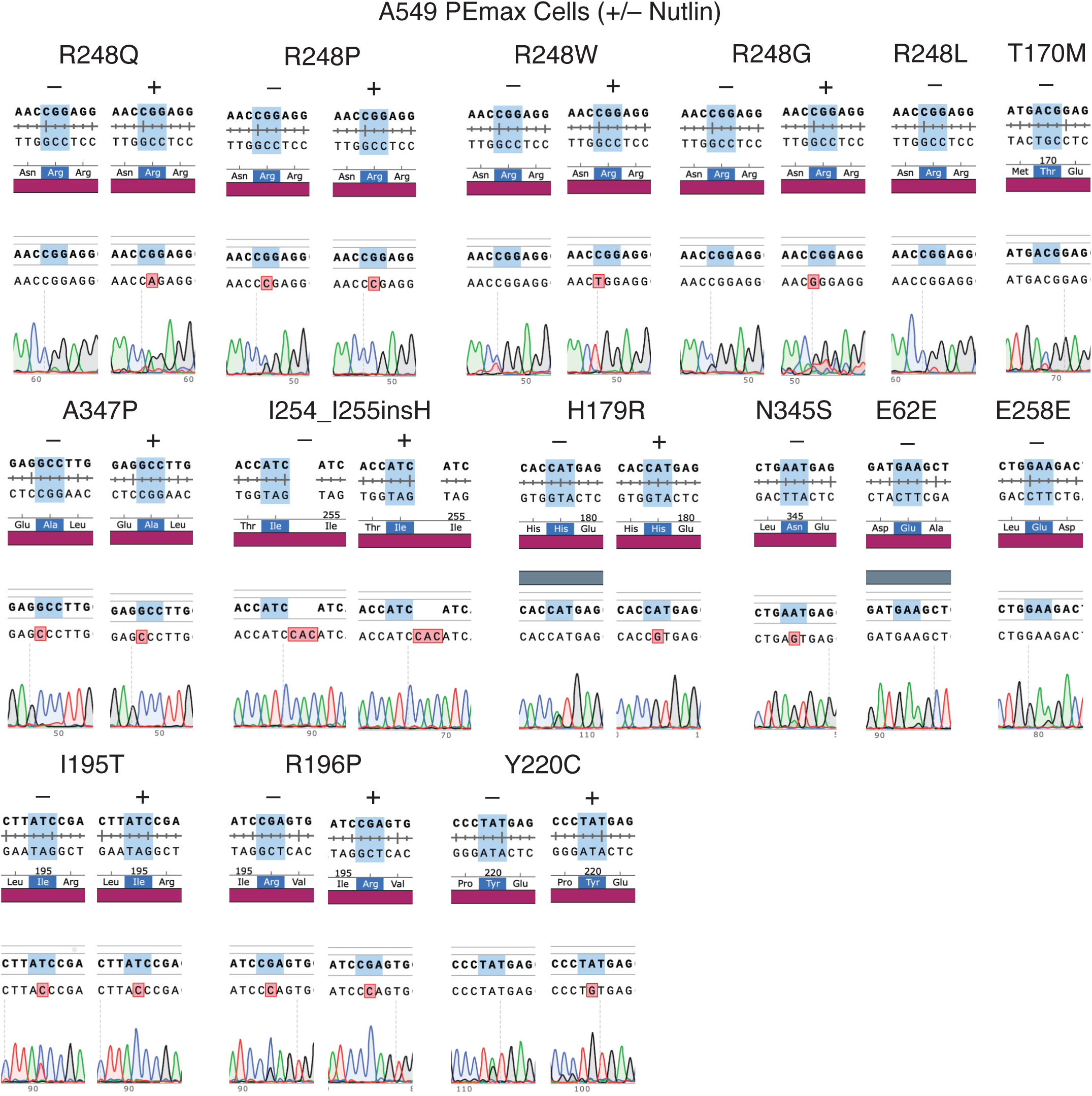
Endogenous sequencing of TP53 validates the on-target editing activity of individually tested pegRNAs in multiple cell types. Sanger sequencing traces of on-target editing of pure A549-PEmax lines in the presence (+) or absence (-) of Nutlin.

**Extended Data Fig. 8.**
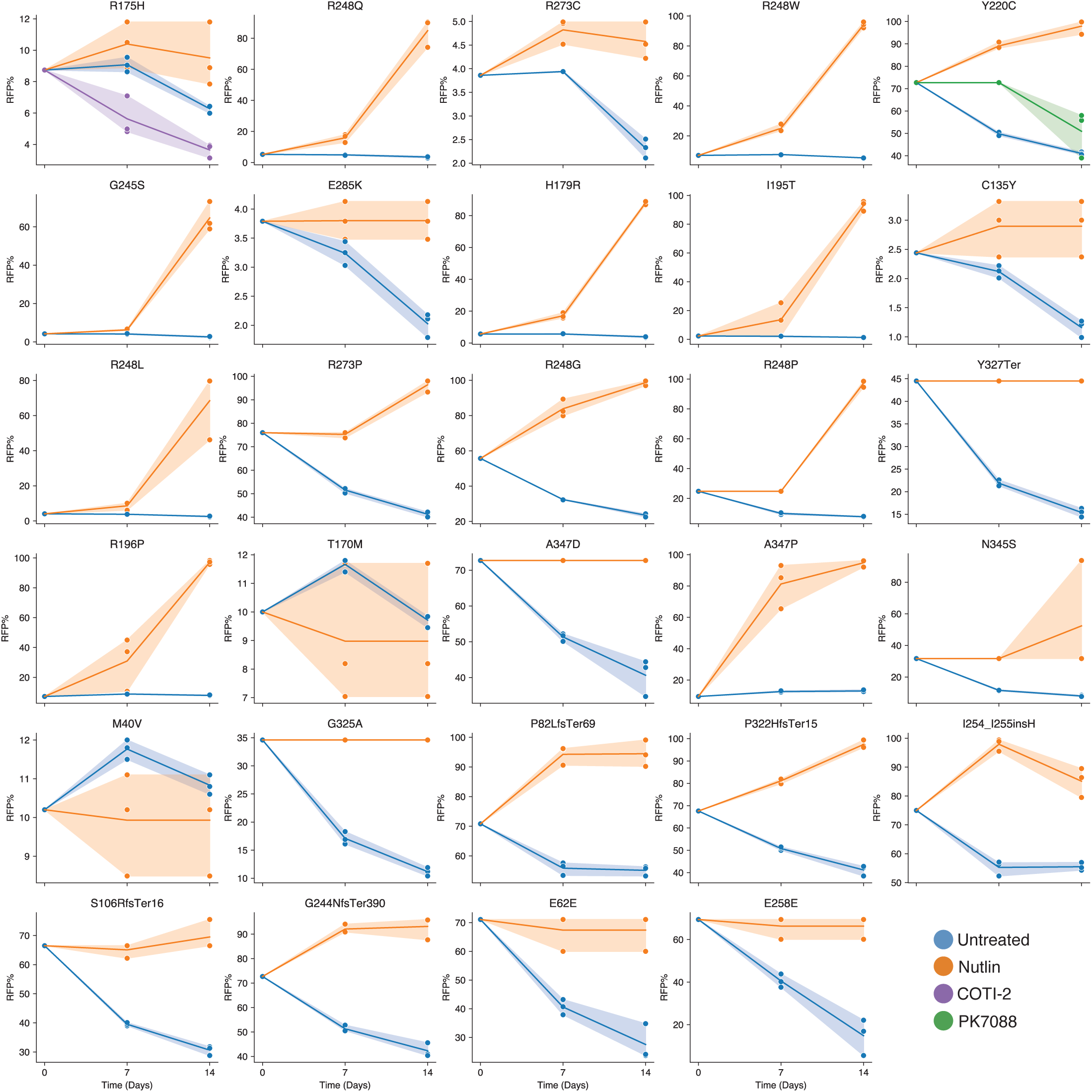
Competition assays functionally validate the pathogenicity of TP53 variants identified with prime editing screens. Full competition assay plots for each of the 29 pegRNAs chosen for validation. Flow cytometric analyses were performed at days 7 and 14 following the initiation of the competition assay. For data points with fewer than 400 flow cytometry events (indicating insufficient viable cells), the RFP positive percentage was set to be equal to that of the previous time-point.

**Extended Data Fig. 9.**
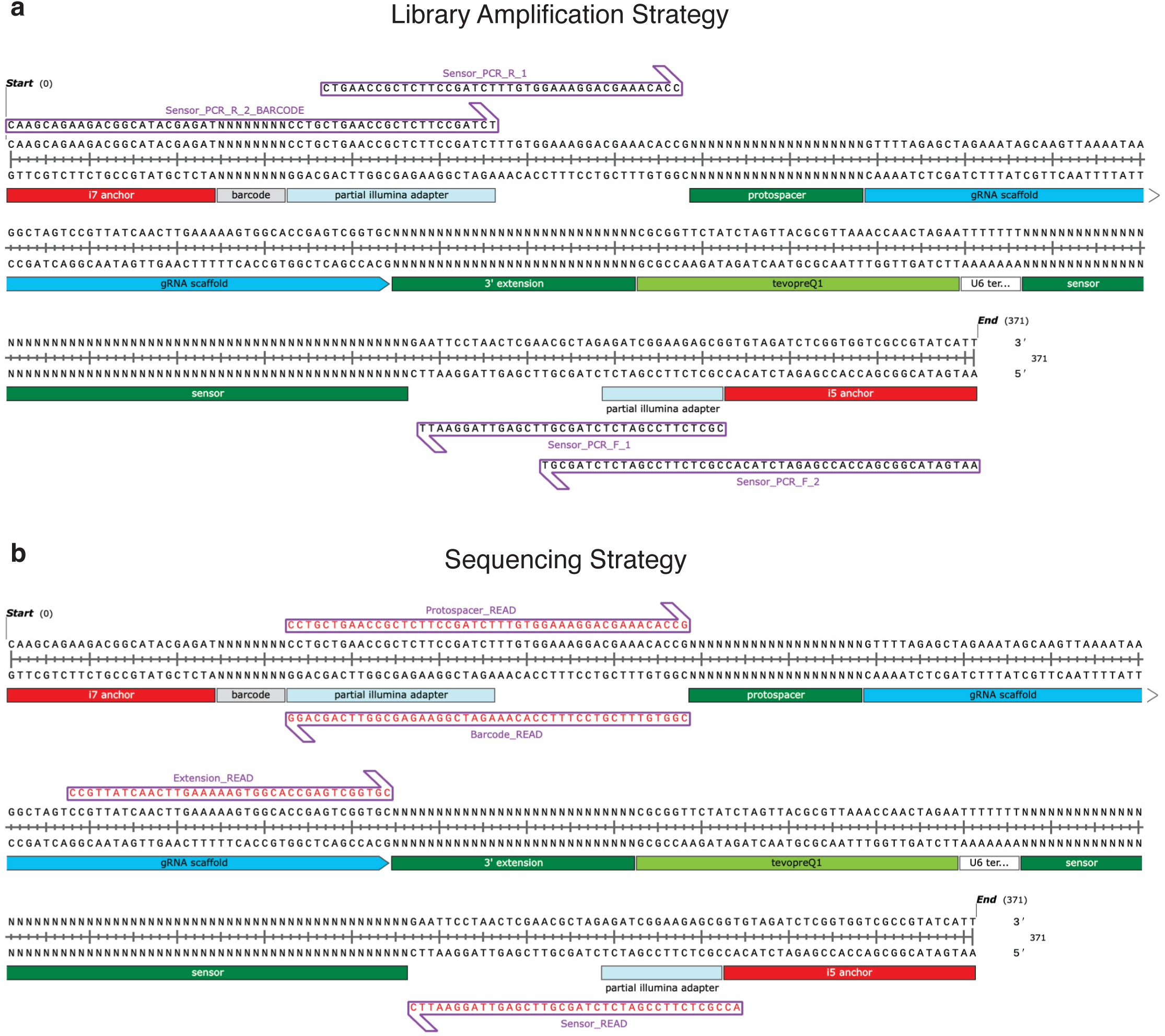
Library amplification and sequencing strategy. **a,** Schematic of the PCR1 and PCR2 strategies used to amplify the pegRNA-sensor cassette for NGS. **b,** Custom sequencing strategy used with the NovaSeq 6000 system. Sensor_Read = Read 1, Extension_Read = Read 2, Barcode_Read = index 1 read, Protospacer_Read = index 2 read.

