## Supplementary material for "High throughput evaluation of genetic variants with prime editing sensor libraries": Gould_et_al_bioRxiv_Sept_2023_Supplemental-Protocol-1.pdf

### PE Sensor CRISPR Library Cloning

#### Workflow:

*Amplification → Insert preparation → Ligation → Electroporation*

#### I - Amplification

##### *Reagents needed:*

- 2X NEBNext Master Mix (#M0541L) (PCR hood room -20C)
- Oligo pool at 1ng/uL (diluted from reconstituted OLS library; see below)
- Sensor_F and Sensor_R primers:

| Sensor_F | Sequence | Sensor_R | Sequence |
|---|---|---|---|
| F | CATAGCGTACACGTCTCACACCG | R | GTGCCGTTGACGACCGGATCTAGAATTC |

##### *Reconstitution of lyophilized OLS library:*

1. Resuspend the pellet in 100 uL of TE Buffer pH 8.0 or QIAGEN EB Buffer.
2. Incubate at RT for ~1hr and periodically vortex to ensure complete resuspension (or put in a shaking block at RT).
3. Nanodrop and aliquot (typically n=10 aliquots per OLS library) and store these at -20 C.
4. If cloning libraries right away, prepare *fresh* serial dilutions until getting a diluted stock at 1 ng/uL (see below).

*Protocol:*

- We typically perform n=4 PCR reactions per pool of ~1000 gRNAs (an excess, but allows for plenty of backup insert for subsequent sub-cloning of libraries into different destination vectors, if desired, or to repeat any that fail QC).
    - In this case, performed 32 parallel PCR rxns (~n=1 PCR per pool of ~1000 pegRNAs)
- All PCR reactions should be set up in a PCR hood following standard procedures.
- Always include a water-only control for every primer set to assess non-specific amplification and/or contamination.

**1. Reaction conditions (50uL reactions): MAKE A MASTERMIX**

- 1 uL of oligo pool* (at 1 ng/uL)
- 1.5 uL of Fwd primer (10 uM stock)
- 1.5 uL of Rev primer (10 uM stock)
- 25 uL of 2X NEBNext
- 21 uL of water
- *The oligo pool should be added outside of the PCR room, preferably in a bench or room that is not used routinely for cloning gRNAs. Never bring oligos or oligo pools into the PCR room.

**2. Cycling conditions**

- 98 C x 30s
- 98 C x 30s
- 53 C x 30s
- 72 C x 30s
- [**GO TO STEP 2 x 18 total cycles]** Go to step 2 x 24 cycles (note: the # of cycles can be varied and has been tested and optimized; anywhere from 10-24 cycles is fine; less is usually better to minimize potential over-amplification but I haven't found this to be an issue; in fact, most libraries I've cloned have been done w/ 24 cycles; if doing 10 cycles, I would run 8 rxns per pool)
- 72 C x 5 min
- 4 C forever
- Named "CRISPR library cloning 25 cycles" under the library cloning folder in the Mastercycler 1 in Bay 1.

Results for optimization of TP53 PE sensor library cycle count (18 cycles chosen):

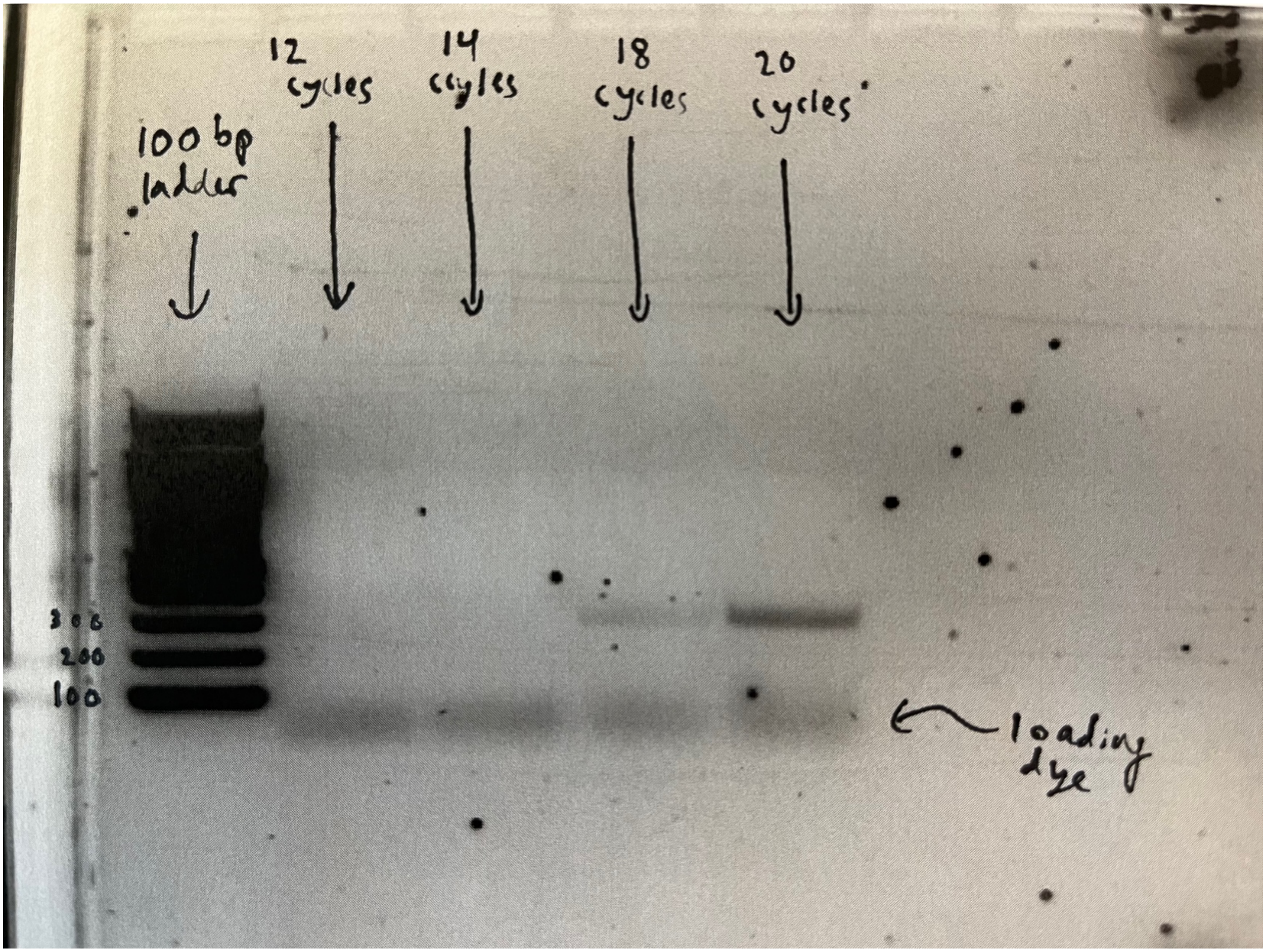


**3. PCR purification**

1. Pool up to four 50 uL reactions per pool and PCR purify using a single QIAGEN column and standard QIAGEN PCR purification protocol.
2. Add 10 uL of 3M NaOAC pH 5.2 for every 5 volumes of PB used per 1 volume of PCR reaction (e.g. 200 uL pooled rxns need 1 mL of PB + 10 uL NaOAC).
3. Elute in 50 uL of pre-warmed (55 C) EB.
4. Run 5 uL of each purification in a gel (it should look like the gel below).

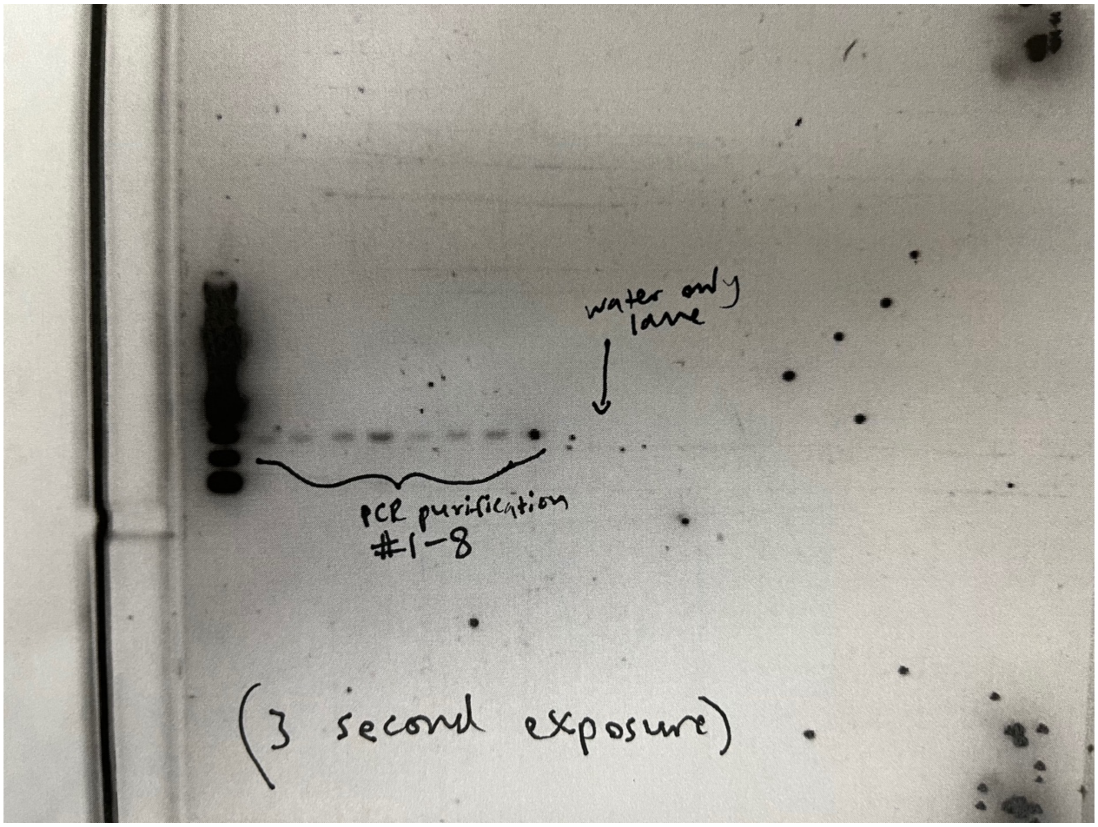


#### II - Insert preparation

- Insert digestion with Esp3I (NEB) and EcoRI-HF (60 uL reactions)
    - 45 uL purified PCR product **(*8 = 360 uL MM)**
    - 6 uL rCutSmart NEB buffer (10X) **(*8 = 48 uL MM)**
    - 3 uL Esp3I NEB **enzyme (*8 = 24 uL MM)**
    - 3 uL EcoRI-HF NEB Enzyme **(*8 = 24 uL MM)**
    - 3 uL water **(*8 = 24 uL MM)**
    - Digest at 37 C x 4 hrs
- Insert purification
    - Pool up to 4 reactions per pool and PCR purify using a single QIAGEN column.
    - Add 10uL of 3M NaOAC pH 5.2 for every 5 volumes of PB used per 1 volume of PCR reaction.
    - Elute in 30 uL of pre-warmed (55C) EB.

#### III - Ligation

1. Ligation to cut backbone (20 uL reactions)

    - **Performing n=16 parallel ligations**
    - 6 uL cut and dephosphorylated backbone (at 50 ng/uL)
    - 3 uL insert (at 1 ng/uL)
    - 2 uL T4 ligase buffer (10X)
    - 1 uL T4 ligase (high concentration; #M0202M; do not use low concentration ligase)
    - 8 uL water
    - Incubate at 16 C overnight
2. Precipitation of ligation reactions
    - Pre-spin PhaseLock tubes at max speed for 5 min.
    - Pool up to 4 ligations per pool and complete to 300 uL using water.
    - Add 300 uL of equilibrated Phenol (no Chloroform or Isoamyl alcohol; ensure you pipet from bottom phase).
    - Mix and extract using PhaseLock tubes (spin at max speed for 5min).
    - After spin, take the top 250 uL watery phase, add 25 uL 3M NaOAC pH 5.2, 750 uL ice cooled EtOH and 1.5 uL Pellet paint (Novagen).
    - Mix and store in -20 C (overnight or longer). Can also do -80C for 2hrs.
    - Spin down (13K RPM for 30 min at 4C).
    - Discard supernatant and add ~1 mL of 70% EtOH.
    - Spin down (13K RPM for 5 min at 4C).
    - Repeat for another 70% EtOH wash.
    - Dry and resuspend in pre-warmed EB (55 C) (3 uL of EB per 4 precipitated reactions).

#### IV - Electroporation

#### Steps:

1. **Bacterial electroporation**
    A. Dry 10 cm/15 cm LB-Amp / LB-Carb plates at 37 C for at least 6 hrs until completely dry.
    B. Pre-chill cuvettes at -20 C / -80 C throughout the day and store at -20C when ready.
    C. Thaw electrocompetent cells (e.g. Lucigen Endura ElectroCompetent Cells; #60242-2) on ice and aliquot 25uL of bacteria per pre-chilled eppendorf tube per transformation (typically 1 transformation per precipitated ligation reaction, and 1 of these for 1 pool of ~1000 gRNAs).
    D. Add 3uL of precipitated ligation to bacteria.
    E. Incubate on ice for 10min.
    F. Transfer bacteria to pre-chilled cuvette (~28uL) (wipe sides of cuvette w/ kim-wipe before electroporating to remove condensation).

G. Electroporate (manual setting, 2.00 kV, aim for at least 5.2 msec).
H. Rapidly quench w/ ~980uL pre-warmed (37C) SOC or LB.
I. Recover at 37C x 1hr in a bacterial shaker.

2. **Plating**
   A. Dilution plates
      1. Set up serial dilution plates (10E2 - 10E6) by taking 10uL of bacteria and diluting in 990uL SOC/LB (initial 10E2 dilution) and then serially dilute (10uL bacteria + 90uL SOC/LB) until obtaining the 10E6 dilution.
      2. Plate 100uL dilutions into 10cm pre-warmed plates (make sure these are dry).
      3. Spread thoroughly using 4 glass beads per plate until dry.
      4. Incubate inverted at 37C overnight for 16hrs.
      5. Count colonies next day; ideal representation = 10,000X.
      6. SEND FOR SEQUENCING
   B. Library plates
      1. Plate ~240uL of bacteria per 15cm plate (divide in 4 spots in the plate).
      2. Spread thoroughly using 4 glass beads per plate until dry.
      3. Incubate inverted at 37C O/N for 16hrs.
3. **Scraping**
      1. Prepare at least 250mL of fresh LB-Amp per four 15cm plates.
      2. Add 20mL of LB-Amp per 15cm plate.
      3. Scrape using cell lifters.
      4. Transfer to 1L flask.
      5. Repeat steps 1-4.
      6. Complete to 250mL with fresh LB-Amp.
      7. Shake at 37C for at least 2hrs (up to 4hrs).
      8. Spin bacteria and freeze pellets or proceed to large-scale maxipreps.
