## Supplementary material for "High throughput evaluation of genetic variants with prime editing sensor libraries": Gould_et_al_bioRxiv_Sept_2023_Supplemental-Protocol-2.pdf

### pegRNA Golden Gate Assembly

#### Workflow:

1. Top/bottom oligo design → (synthetic Golden gate assembly in snapgene) → phosphorylation/annealing of oligos → oligo dilution 1:100 → Golden Gate reaction → transformation/plating

This is a general schematic, though with BsaI, rather than BsmBI as the Golden Gate cutter (thus overhangs aren't accurate):

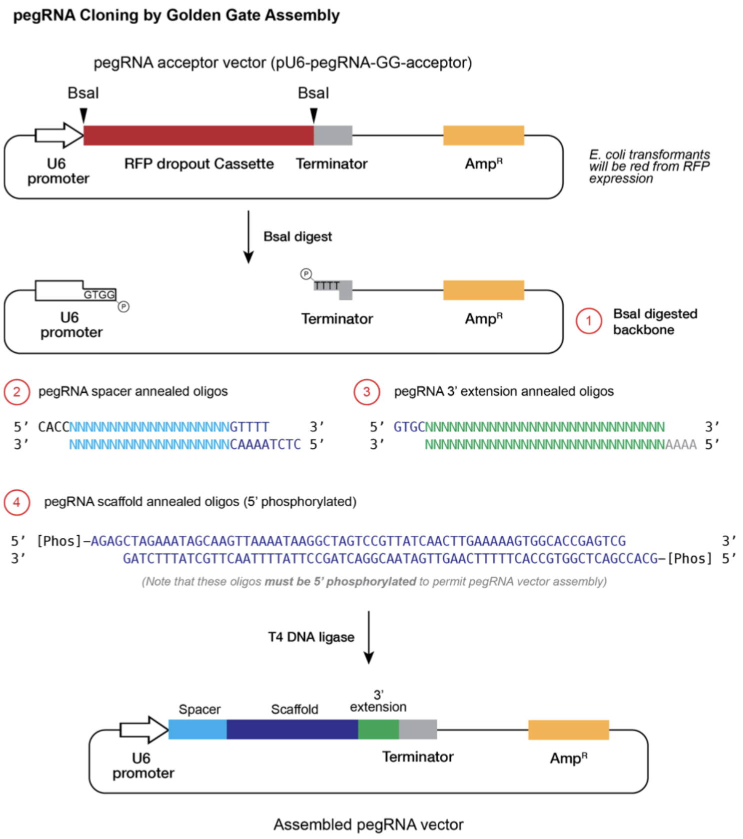


#### Oligo Design:

**Protospacer top/bottom: (5' → 3' and 3' → 5'):** **<u>(Make sure protospacer starts with G)</u>**

CACC(G)NNNNNNNNNNNNNNNNNNNGTTTT

(C)NNNNNNNNNNNNNNNNNNNCAAAATCTC

**Scaffold top/bottom: (5' → 3' and 3' → 5'):**

AGAGCTAGAAATAGCAAGTTAAAATAAGGCTAGTCCGTTATCAACTTGAAAAAGTGGCACCGAGTCG

GATCTT TATCGTTCAATTTTATTCCGATCAGGCAATAGTTGAACT TTTTCACCGTGGCTCAGCCACG

*Last modified by SIG - 6/2023*

**3’ Extension top/bottom: (5’ → 3’ and 3’ → 5’):**

GTGCNNNNNNNNNNNNNNNNNNNNNCGCGGTTCTATCTAGTTACGCGTTAAACCAACTAGAA
NNNNNNNNNNNNNNNNNNNNNGCGCCAAGATAGATCAATGCGCAATTTGGTTGATCTTAAAA

Modified version with tevopreQ1 included in backbone:

GTGCNNNNNNNNNNNNNNNNNNNNN
NNNNNNNNNNNNNNNNNNNNNGCGC

**Key: Overhang, Scaffold Region, tevopreQ1 motif**

#### Materials needed:

| Materials/Equipment | Location | ☐ |
|---|---|---|
| Top/bottom oligos @ 100 uM | N/A | |
| T4 DNA Ligase | Bay 2 | ☐ |
| T4 PNK | -20C (near Bay 8) | ☐ |
| 10X T4 DNA Ligase Buffer | -20C (near Bay 8) | ☐ |
| Nuclease-free (NF) water | Bay 1, 2 | ☐ |
| NEB Golden Gate Enzyme Mix **(BsmBI)** | -20C (near Bay 8) | ☐ |
| UPEmS or UPEmS_tevo (or other vector) @75 ng/uL | -20C | ☐ |

#### Steps:

1. **Run annealing and phosphorylation reactions for oligos**
   a. There are 3 oligo pairs: (1) protospacer top/bottom, (2) scaffold top/bottom, (3) 3’ extension top/bottom. These oligos should be **generated with the appropriate overhangs** according to the map above. ***Performing an in silico Golden Gate reaction in SnapGene is HIGHLY recommended before proceeding.***
   b. Phosphorylate and anneal the **scaffold** oligos. Alternatively, we generally keep a supply of phosphorylated scaffold in stock that can be re-used:

| Component | 1x |
|---|---|
| **Oligo 1 (100 µM)** | **1uL** |
| **Oligo 2 (100 µM)** | **1uL** |

| | |
|---|---|
| 10x T4 DNA ligase buffer | 1uL |
| $H_2O$ | 6.5uL |
| T4 PNK | 0.5uL |
| Run annealing program | |

c. Anneal (but do not phosphorylate) the **protospacer and 3' extension oligos** (same reaction as above, but with T4 DNA Ligase instead of T4 PNK):

| Component | 1x |
|---|---|
| Oligo 1 (100 µM) | 1uL |
| Oligo 2 (100 µM) | 1uL |
| 10x T4 DNA ligase buffer | 1uL |
| $H_2O$ | 6.5uL |
| T4 DNA Ligase | 0.5uL |
| Run annealing program | |

**Annealing program:**

| | |
|---|---|
| 37˚ C | 30 mins |
| 95˚ C | 5 mins |
| 25˚C FINAL | **Ramp down to 25˚C at 5˚C/min** (i.e. 0.083˚C/sec; 0.1˚C/sec also seems to work fine) |

**2. Dilute phosphorylated and annealed oligos 1:100 in nuclease-free water.**

a. Keep products on ice and store at -20˚C.

##### 3. Perform Golden Gate reaction

| Component | 1x | 12x | 5x |
|---|---|---|---|
| UPEmS Vector or other (75 ng/µL) | 1.5µL | 18 | 7.5 |
| Protospacer (100nM) | 1µL | 12 | 5 |
| scaffold (100nM) | 1µL | 12 | 5 |
| 3' ext (100nM) | 1µL | 12µL | 5µL |
| T4 DNA Ligase Buffer (10x) | 2uL | 24 | 10 |
| NEB Golden Gate Enzyme Mix | 1uL | 12 | 5 |
| $H_2O$ | 12.5µL | 150 | 62.5 |
| TOTAL | 20uL | 240uL | 100 uL |
| **(42C, 1min → 16C, 1min) x 60 repeats → 60C, 5min** | | | |
| **Transform and plate the entire 20 uL reaction** | | | |

##### 4. Transformation reaction copied for convenience:

#### Materials needed:

| Materials/Equipment | Location | ☑ |
|---|---|---|
| Ice | Hallway (near TC) | ☐ |
| Competent cells (very fragile!) | -80 C | ☐ |
| Plasmid | -20 C (near Bay 8) | ☐ |
| Block for heat shock @ 42 C | Bay 1 in Lab | ☐ |
| SOC/LB | Hallway (near the lab entrance) | ☐ |
| Petri dishes | Hallway (near the lab entrance) | ☐ |
| Glass beads | Bay 2 and 3 in Lab | ☐ |

#### Steps:

1. Always thaw competent cells on ice as they are quite fragile.
2. Check the tube. Are competent cells at the bottom of the tube? If not, give it the "manual centrifuge" treatment by giving the tube a quick outward whip to bring them to the bottom of the tube.

3. Add **20 uL of the golden gate reaction product** to a vial of competent cells and gently flick the tube. Ensure bacteria are at the bottom of the tube.
4. Incubate on ice for 5-10 min
5. Heat shock at 42 C for 30-45 seconds.
6. Incubate on ice for 3min.
7. Add 250uL SOC or LB-only media. At this point, the bacteria can sit at room temperature for up to 1hr.
8. Plate or inoculate plasmid prepping cultures.
9. Plate 50-100uL bacteria and thoroughly spread with glass beads.
10. Incubate inverted plates at 37C overnight.
